# Transient morphogenetic constraints organize self-assembling axon neighborhoods for robust yet flexible wiring

**DOI:** 10.64898/2026.09.09.750528

**Authors:** Christopher A. Brittin, Anthony Santella, Kristopher Barnes, Mark W. Moyle, Li Fan, Ryan Christensen, Irina Kolotuev, William A. Mohler, Hari Shroff, Daniel A. Colón-Ramos, Zhirong Bao

**Affiliations:** Developmental Biology Program, Memorial Sloan Kettering Cancer Center, New York, NY 10065, USA; Molecular Cytology Core, Memorial Sloan Kettering Cancer Center, New York, NY 10065, USA; Graduate Program in Neuroscience, Weill Cornell Medicine, New York, NY 10065, USA; Departments of Neuroscience and Cell Biology, Yale University School of Medicine, New Haven, CT 06536, USA; Janelia Research Campus, Howard Hughes Medical Institute, Ashburn, VA, USA; Department of Biomedical Sciences, Faculty of Biology and Medicine, University of Lausanne, Lausanne, Switzerland; Center for Integrative Genomics, University of Lausanne, Lausanne, Switzerland; Department of Genetics and Genome Sciences and Center for Cell Analysis and Modeling, University of Connecticut Health Center, Farmington, CT 06030, USA

## Abstract

Nervous systems form wiring patterns that are reproducible across individuals. This reproducibility is thought to emerge from molecular encoding and developmental events, but their relative contributions remain unclear. We address this question in the *C. elegans* neuropil, where embryonic developmental dynamics and adult anatomy are resolved at single-cell resolution. We find that transient morphogenetic structures — rosettes, corridor cells, pioneer axon scaffold — restrict which axons make contact, shaping the neuropil into overlapping neighborhoods. This demonstrates how early events constrain wiring choices, but not whether they explain the resulting reproducibility. To explain, we use an agent-based model of stochastic innervation that recapitulates macro- and micro-level reproducibility, revealing a trade-off between physical constraint and molecular specificity that limits neighborhood size. Counterintuitively, less selective axons produce more reproducible wiring when constrained within neighborhoods. This trade-off lets nervous systems maximize reproducibility without having to molecularly encode every axon-contact, a strategy for robust yet flexible wiring.

## Introduction

Nervous systems form elaborate wiring patterns that are reproducible across individuals. Broadly, two accounts have emerged to explain this reproducibility^1^. One emphasizes that combinations of surface recognition molecules specify how individual neurons wire together (e.g. DSCAMs and protocadherins)^2,3^. However, decades of research have not established this hypothesis as a general principle, prompting calls to move beyond molecular codes^4^. This second account views wiring as a morphogenesis problem^4,5^ where cell movements and transient tissue structures restrict axon encounters^6^, constraining where molecular interactions can operate. These accounts are not mutually exclusive, but together raise the question of how much molecular specificity is required once morphogenetic constraints have been enacted.

Technical advances now sharpen this question^7^, letting us quantify wiring reproducibility across individuals with single-axon resolution^8^. This reveals that reproducibility is not uniform: some axon-contacts are consistently reproduced across individuals while others are not^9^. Studies have found that this pattern is structured rather than random^10^, suggesting that reproducibility itself carries developmental significance, rather than reflecting mere noise^10,11^. Across a population, wiring is then best described by a distribution of reproducibility rather than by any single canonical pattern. An integrated account of molecular specificity and morphogenetic constraint must explain this full distribution.

The *C. elegans* main neuropil (nerve ring) provides a tractable system to build such a model. Because individuals are genetically identical^12^, differences in wiring patterns must reflect stochastic development rather than genotype. Nerve ring wiring reproducibility has been characterized at two scales. At the macro-scale, spatially and compositionally stereotyped domains emerge from population-conserved axon-contacts^13,14^, creating regions where axon groups preferentially interact. At the micro-scale, pairwise axon-contacts are not uniformly conserved across individuals^9,13^: some contacts are reproducible across the population while others occur only in subsets of individuals, producing a bimodal distribution of contact reproducibility^13^. Despite comprehensive single-cell mapping of lineage, position, connectivity and molecular identity across life-stages^9,15–20^, no developmental model has been offered to account for wiring reproducibility at either scale.

This study combines embryonic imaging, molecular analysis, and computational modeling to ask how molecular encoding and early developmental events jointly modulate wiring pattern reproducibility in the nerve ring. We focus on how the embryonic microenvironment physically constrains the packing and sorting of axons within the nascent neuropil (axon patterning^21^). We trace the formation of transient multicellular rosettes that define nerve ring innervation entry points, and identify corridor cells that constrain pioneer fascicle trajectories. These pioneers self-organize into a scaffold that establishes spatial domains, and we identify molecular expression patterns that would let follower axons distinguish among pioneers as they innervate. These observations demonstrate how early events constrain wiring patterns, but they do not directly explain wiring pattern reproducibility. For this, we built an agent-based model that simulates nerve ring innervation while varying physical constraints and molecular specificity. When fit to the nerve ring’s domain and bimodal reproducibility patterns, the model shows that these two factors interact to restrict axon innervation to local neighborhoods which increases wiring reproducibility (Fig. 1).

**Figure 1:**
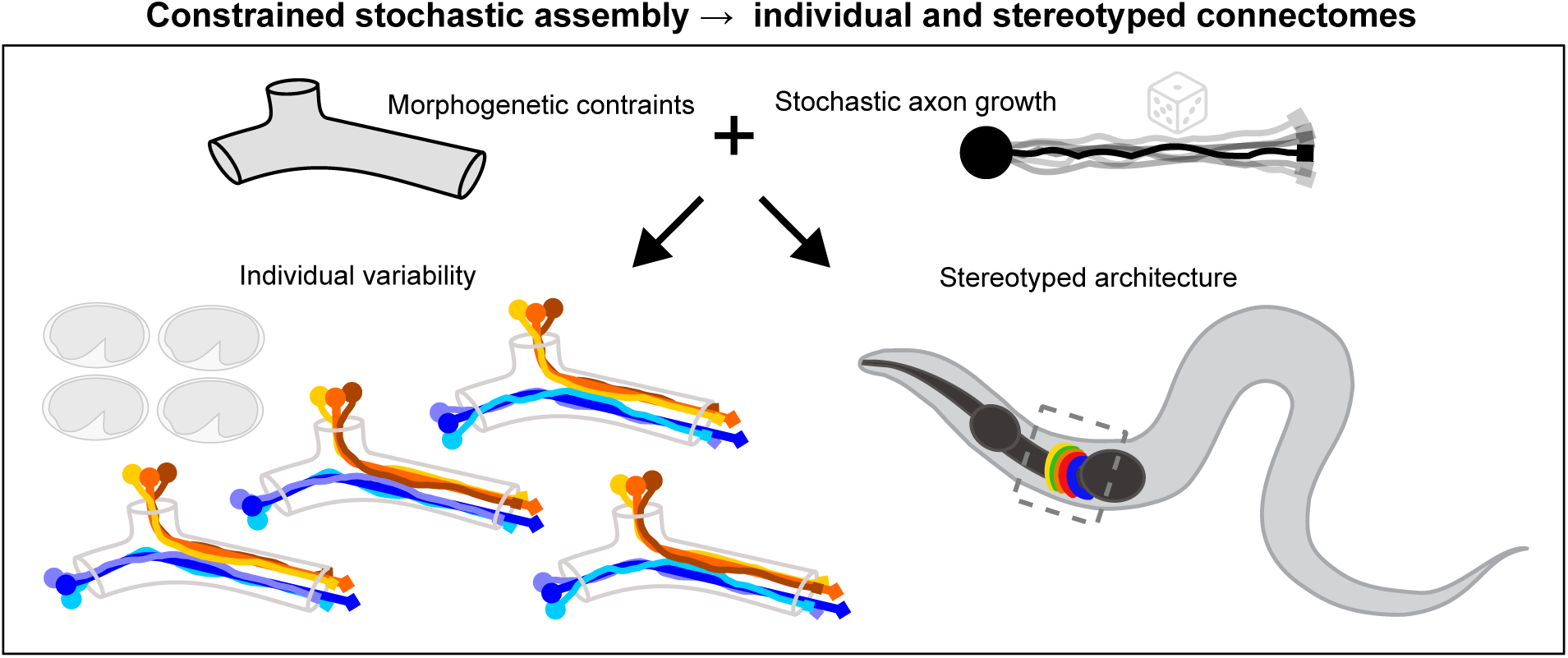
Morphogenetic constraints enable reproducible circuit assembly from stochastic axon growth. Conceptual overview of constrained stochastic assembly. During development, transient morphogenetic constraints define local neighborhoods that restrict which axons can encounter one another, raising the question of how much molecular specificity is required once these constraints are enacted. Stochastic growth within these neighborhoods, governed by a trade-off between physical constraint and molecular specificity, produces individual-specific wiring while yielding a conserved global architecture.

### Rosettes define discrete entry points for nerve ring assembly

To identify the emergence of early tissue structure, we used time-lapse imaging to track single-cell lineages (PIE-1::mCherry) and cell morphology (UNC-33::GFP) during the earliest stages of nerve ring formation (Table S1). At ∼350 minutes post fertilization (mpf), eight multicellular rosettes appeared in a bilaterally symmetric semicircle around the pharynx (Fig. 2a). Rosettes form as multiple cells constrict toward a central vertex^22^ and were detected via membrane convergence (Fig. 2b,c). Their cellular composition was verified by lineage tracing from early embryonic cleavage (Fig. 2d).

**Figure 2:**
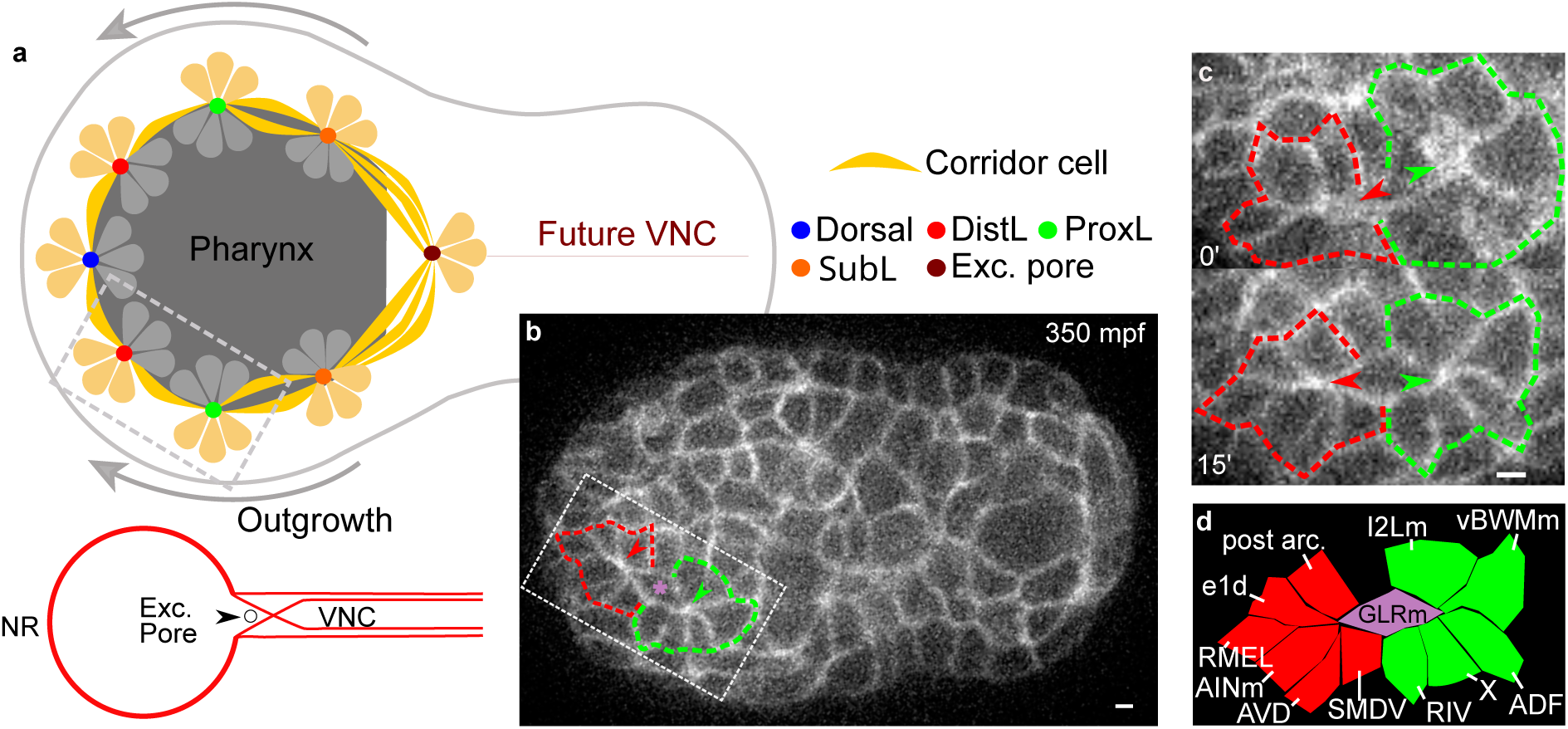
Early morphogenesis defines entry geometry for nerve ring innervation. **a**, Top: schematic of 8 rosettes prior to nerve ring innervation. Thin black line: embryo contour. Ventral view, anterior to left. Color dots are rosette centers. Thick arrows: direction of axon outgrowth. Cell composition of rosettes in Table S2. Bottom: schematic of final central nervous system. The posterior side of the excretory pore marks the site where axons exit the nerve ring to enter the ventral nerve cord (VNC). **b**, Embryo (350 mpf) with UNC-33::GFP expressed in membrane. Ventral view, anterior to left. Grey box corresponds to grey box in **c**. * marks corridor cell between DistL and ProxL rosettes. **c**, Convergence of rosettes in **b** over 15 minutes. **b**, **c**, Dashed line and arrowheads mark the contour and center of DistL (red) and ProxL (green) rosettes, respectively. Identical histogram adjustments applied to all images to improve contrast on dark background. Scale bar: 1 μm. **d**, Identities for rosette cells in **c**. Purple: corridor cell. Cell identities were assigned by lineage tracing during live imaging. X marks cell ABaxxxxx which subsequently undergoes programmed cell death.

Rosette formation is spatiotemporally stereotyped relative to internal landmarks (Fig. S1, Movie S1). The dorsal rosette arose at the anterior midline, while the excretory pore rosette formed posteriorly, incorporating neurons from both the nerve ring and ventral cord. Between these poles, bilaterally symmetric Distal Lateral (DistL), Proximal Lateral (ProxL), and Sublateral (SubL) rosettes assembled. For SubL, three smaller precursor rosettes merged into a larger 27-cell rosette (Fig. S1a, S2d).

Rosette cell compositions are stereotyped (Fig. 2d, S2; Table S2,‘rosette_composition’). Each rosette includes both neuronal and non-neuronal cells: pharyngeal cells, muscle, glia-like GLRs, or excretory pore cells. The non-neural cells have previously been implicated in supporting the adult nerve ring^23^, suggesting rosettes seed the nerve ring with its structural components. Thus, at the tissue level, rosettes are not just transient morphogenetic structures but the first distributed organizers that embed both neural and non-neural elements into the nascent nerve ring scaffold (Movie S2), thereby defining the space where axons will interact.

### Rosettes support pioneer fascicle orientation toward the nerve ring

Because many rosette cells are neuronal, we used live imaging to dissect how they initialize nerve ring formation. All eight rosettes exhibit a stereotyped sequence: membrane constriction brings cells into contact, fluorescence intensifies at the central vertex, and pioneer axon-like processes emerge from this vertex (Fig. 3a, Movie S3). The SubL rosette generates the first fascicle, including known pioneers SIAD, SIBV, and SMDD^14,24,25^. In the DistL rosette, AVD and AVJ extend in close succession (∼10 min apart). In contrast, some rosette neurons (e.g. papillary sensory cells) delay projection, indicating that the rosette center orients outgrowth but does not induce fascicle extension.

**Figure 3:**
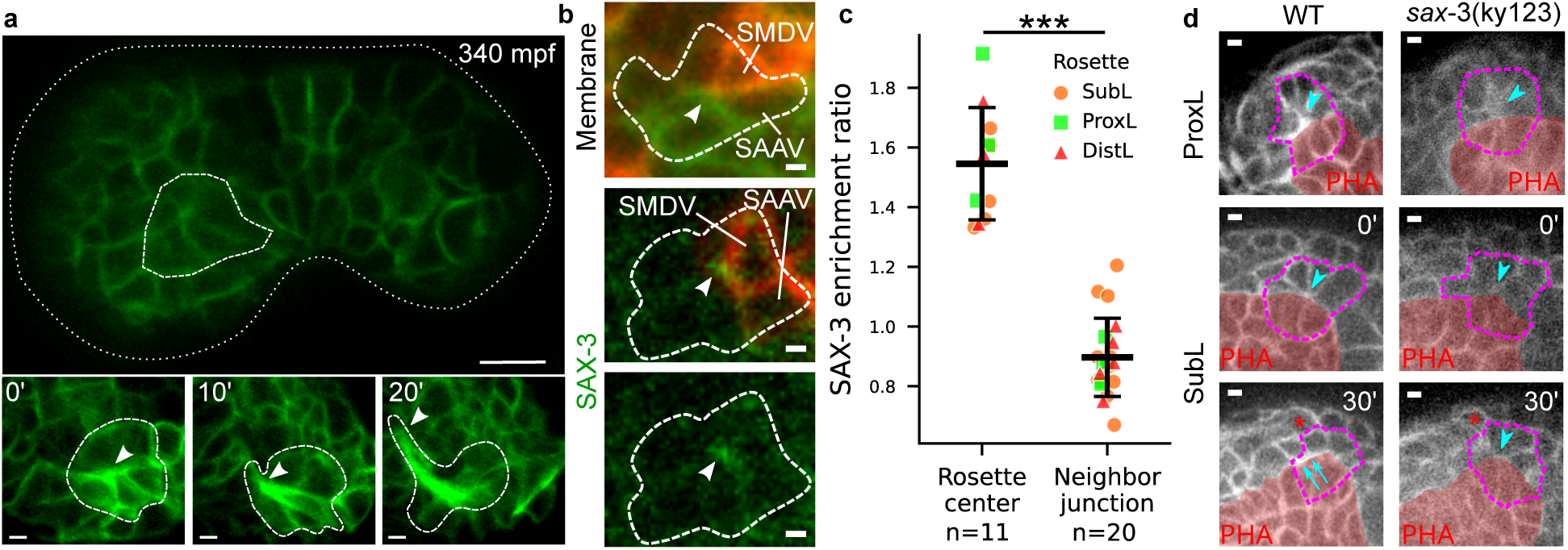
Rosette centers bias and coordinate pioneer axon outgrowth. **a**, Top: SubL rosette (thick dashed line) in UNC-33:GFP (membrane) labelled ∼340 mpf embryo (thin dashed line for contour). Scale bar: 10 μm. Bottom: Select time points of pioneer outgrowth starting at ∼345 mpf. Arrows: rosette center (0 min) and leading edge of growing axons (10 and 20 minutes). Scale bar: 1 μm. **b**, Top: Relative position of CND-1::PH::mCherry-labelled cells (red) within ProxL rosette (dashed line) identified using UNC-33::GFP (green membrane). Arrowhead: rosette center. Middle: SAX-3::GFP localization (arrowhead) relative to mCherry cells. Named cells are ProxL mCherry labelled cells used as landmarks. Bottom: SAX-3::GFP channel alone. Scale bar: 1 μm. **c**, Quantification of SAX-3::GFP enrichment ratio at rosette centers and neighboring cell-cell junctions across SubL, ProxL, and DistL rosettes. Dots indicate individual rosette centers (*n*=11) or junctions (*n*=21); dot color and shape indicate rosette type (SubL: orange circles; ProxL: green squares; DistL: red triangles). Black bar: mean; error bars: 1 s.d. ***: *p* = 3.01×10^−8^ by Welch’s t-test. **d**, Phenotypes in *sax-3*(ky123) mutants. Membrane-expressing UNC-33::GFP images. Top: A ProxL rosette (purple dash) with less focused center (arrowhead) and displaced from pharynx (PHA, red patch) as compared to the wild type (WT). Bottom: The SubL rosette forms as in WT (0 min, arrowheads point to rosette centers). In WT embryos, the SubL rosette converges and generates axon fascicles (30 min, arrows point to thickened membrane signal of axons along the pharyngeal boundary), whereas in *sax-3*(ky123) the rosette remains isolated and fails to initiate similar fascicle outgrowth (30 min, arrowhead). Red * indicates the position of a cell used as a fiducial marker across embryos. Scale bar: 1 μm.

To identify what confers this orienting property, we screened for receptors implicated in axon guidance. Previously implicated in both *C. elegans* axon guidance and patterning^26–28^, SAX-3/Robo was consistently enriched at rosette centers as discrete puncta rather than diffusely along membranes (Fig. 3b,c, Fig. S3a-c). Therefore, we reasoned that *sax-3*(ky123) null mutants could test if rosettes are required for fascicle orientation.

Across 21 embryos, *sax-3*(ky123) null mutant embryos showed incompletely penetrant defects in rosette assembly and fascicle outgrowth (Table S2, ‘SAX-3_phenotypes’). Phenotypic severity ranged broadly: from effectively wild type (*n*=9, Fig. S3d) to severe head morphology defects (*n*=3, not scored, Fig. S3g). Among milder phenotypes, rosettes can fail to converge to a central vertex or become unstable after convergence, with cells detaching and re-joining (Fig. 3d). Fascicle formation was never observed in the absence of rosette convergence (*n*=6, Fig. S3e), indicating rosettes are important for pioneer fascicle orientation. However, rosette convergence alone was not always sufficient for fascicle orientation (*n*=3, Fig. S3f). Fascicle loss occurred at variable positions along the nerve ring path, differing not only between embryos (Fig. S3e,f) but also between the left and right sides of individual embryos (Fig. S3e). This pattern is consistent with either SAX-3 acting on a stochastic developmental process at the individual level, or SAX-3 acting in a complex with cooperating guidance receptors (e.g. UNC-40/DCC^29^); our data do not distinguish between these possibilities.

These observations indicate that rosette integrity and fascicle orientation are experimentally separable: fascicle formation was never observed without rosette convergence, yet convergence alone did not guarantee correct orientation. While we cannot speak to SAX-3’s precise molecular role, the decoupling revealed by *sax-3* loss-of-function indicates that additional factors — stochastic or otherwise — shape fascicle orientation independently of rosette integrity.

### Corridor cells spatially constrain pioneer trajectories along the nerve ring

We asked how fascicle outgrowth is coordinated across rosettes. Substrates guide pioneer axons in other systems^30,31^. Similarly, our live imaging shows that adjacent rosettes are linked by 1–3 “corridor” cells that span the nerve ring path (Fig. 2b; Table S2, ‘rosette_composition’). As further corroboration, our 340 mpf embryo EM (electron microscopy) reconstruction shows the SubL and ProxL rosettes engaged by the IL2V and IL1Vm mother cells while SIAD, SMDD, and SIBV extended nascent axons from the SubL rosette vertex along these corridor cells (Fig. 4a). These data suggest the corridor substrate guides pioneer extension.

**Figure 4:**
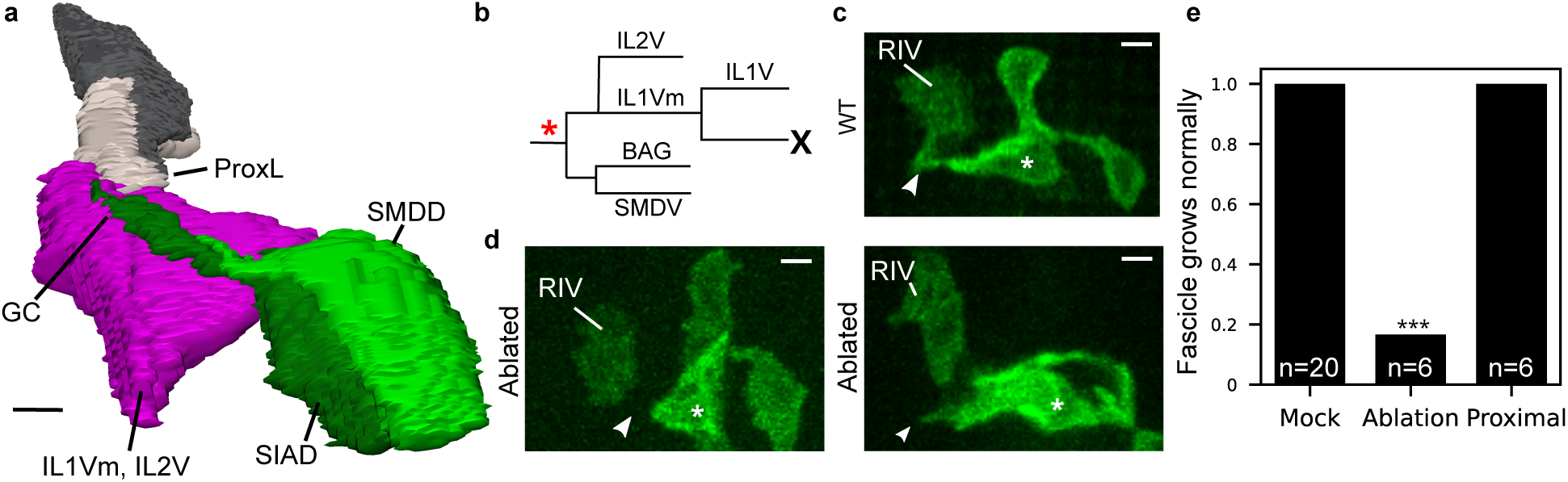
Corridor cells restrict pioneer trajectories during nerve ring assembly. **a**, Volumetric reconstruction from serial-section EM shows initial SIAD (dark green) and SMDD (light green) outgrowth from the SubL rosettes along the corridor cells (IL1Vm and IL2, magenta), directing axons towards the ProxL rosette (gray cells on left). Scale bar: 1 μm. GC: SIAD growth cone. **b**, Target cell for laser ablation (*) and its sublineage. **c**, SubL and ProxL neurons with *lim*4p::PH::GFP label. Arrowheads point to axon tips from SubL neurons. * labels SIAD. In WT, the outgrowth converges with RIV from the ProxL rosette. **d**, Embryo after laser ablation, illustrating the characteristic stall-then-wander sequence: SubL outgrowth initially stalls and fails to reach the ProxL rosette (Left), before extending aberrantly rather than correctly at a later time point (Right, same embryo after a 45° rotation along the anterior-posterior axis, showing the aberrant anterior outgrowth of SIAD more clearly). Full timepoint series confirming both the stall-then-wander sequence and RIV extension (not visible in this focal plane) across all successful ablations are shown in Fig. S4c. Per-embryo outcomes are listed in Table S2 (‘Corridor_ablation’). Scale bar: 1 μm. **e**, Fraction of proper SubL outgrowth under different conditions. Mock: no ablation. Proximal: a cell adjacent to the intended target in **b** was ablated, to control for potential off-target effects. *** denotes a significant difference from mock controls (*p* = 5.8 × 10^−8^, binomial test). *n*: number of embryos.

To test this, we laser ablated the grandmother precursor of corridor cells’ IL2V and IL1Vm (Fig. 4b), which yielded a stall-then-wander phenotype (Fig. 4c,d; Fig. S4a,b; Movie S4). In 5 of 6 embryos with clean ablations, SubL neurons exhibited a characteristic stall following corridor cell removal before eventually extending (Fig. S4c). Of the stalled embryos, 3 subsequently showed wander with incorrect outgrowth and 1 recovered correct outgrowth (Fig. S4f; remaining embryo was inconclusive). Ablation did not affect rosette formation, indicating that the defect was specific to extension along the ring path. Ablation significantly reduced the fraction of embryos with correct outgrowth relative to both mock-ablated and proximal-control groups (Fig. 4e). Control ablations of nearby cells did not perturb normal growth (Fig. S4e), confirming that the phenotype is specific to corridor cell loss. Although ablations also removed BAG and SMDV (Fig. 4b), neither directly contacted SubL outgrowths.

Together these observations indicate that corridor cells guide pioneers along the correct trajectory promptly and without misdirection. Corridor cells transform isolated rosettes into a coordinated chain, restricting pioneers to a common trajectory and preventing early divergence that would expand the axon–axon interacting space, though whether this substrate is passive or contributes active signaling remains unresolved.

### Pioneer fascicles assemble a spatial scaffold that prefigures adult domains

To determine how pioneer axons establish a scaffold for nerve ring assembly, we combined EM^32^ with live imaging of pioneer-specific markers. The 340 mpf EM embryo shows SIAD, SMDD, and SIBV projecting from the SubL rosette (Fig. 4a), confirming their pioneer role^14,24,25^. In a 365 mpf EM embryo, 14 bilateral axons originating from rosettes reached the dorsal midline of the ring (Fig. S5e). Fluorescent markers for 12 of these axons (Dorsal: *ceh-10*; DistL: *zag-1*; ProxL: *cnd1*; ProxL/SubL: *lim-4*; Table S2, ‘pioneer_markers’) reveal a stereotyped, sequential outgrowth, starting with the SubL rosette (Fig. 5a,b, Fig. S5; Table S2, ‘pioneer_triangulation’). Ablation of most pioneer axons disrupts nerve ring formation^14^, while selective SubL pioneer ablation leaves ProxL fascicles intact^33^, showing independent pioneer contributions.

**Figure 5:**
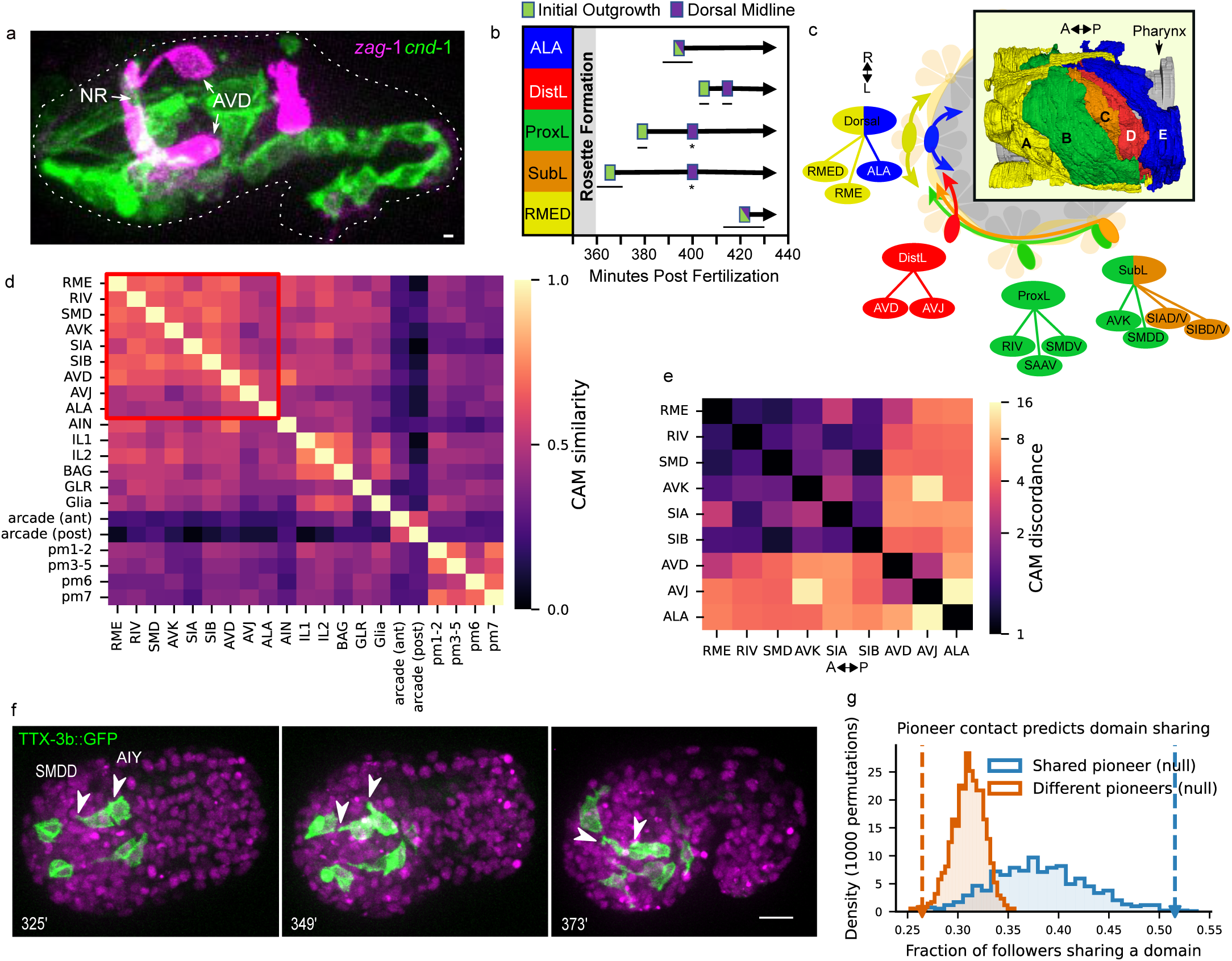
Pioneer fascicles self-sort into a spatial scaffold that prefigures domains. **a**, Example embryo (450 mpf) with *zag-1* and *cnd-1* markers used to track outgrowth dynamics. Dashed white line: contour of the embryo. **b**, Temporal dynamics of axon outgrowth. Horizontal bars show duration of outgrowth (*n*=3 embryos per bar). Timing across embryos is synchronized to when the bilateral ProxL axons meet at the dorsal midline (). **c**, Schematic of how rosette convergence and pioneer sorting give rise to nerve ring spatial organization. Box: Left lateral view of a larval EM-reconstructed nerve ring. Axon colors indicate spatial domain (Table S4); 5–10 axons shown per domain for clarity. Underlay: schematic of pioneer fascicle outgrowth along the rosette chain (L ↕ R), grouped by rosette and colored by domain assignment. **d**, Heatmap of pairwise CAM correlation among pioneers (red box) and surrounding corridor-cell tissue, computed across all annotated CAM genes (Methods). Pioneers show higher mean pairwise correlation with one another (*r* ≈ 0.57) than with corridor cells (*r* ≈ 0.4), indicating pioneers are molecularly distinguishable from surrounding tissue. **e**, Heatmap of pairwise CAM discordance between pioneers, restricted to the 50 most variable CAMs (Methods). Rows/columns: pioneer cells, ordered by anterior-posterior axon placement. **f**, Timelapse of initial AIY and SMDD outgrowth (TTX-3b::GFP, cell-specific; PIE-1::mCherry, pan-nuclear). AIY’s growth cone projects variably at 16 min, then innervates along SMDD by 32 min. **g**, Permutation test relating pioneer contact to domain membership in the larval EM network. Observed domain-sharing frequency among followers: 0.51 (shared pioneer) vs. 0.26 (different pioneers). Permuted null: 0.38 ± 0.046, 0.31 ± 0.015. Solid lines: null distribution from 1000 permutations of pioneer-contact assignments. Dashed lines and arrowheads: observed values. Blue: followers sharing a pioneer; orange: followers of different pioneers. **a**,**f**, Scale bar: 10 μm. **c**,**e**, A ↔ P: anterior-posterior axis.

Pioneer fascicles prefigure the adult nerve ring spatial domains (Table S2, ‘pioneer_domains’). SubL, ProxL, and DistL pioneers map to domains C, B, and D; cells in the Dorsal rosette separate to form anterior and posterior boundaries of the neuropil (RME/RMED → domain A, ALA → domain E). When bilateral axons meet at the dorsal midline, all domains are represented, and fascicle positions align with future domain order (Fig. 5c, Fig. S5d-e).

To serve as a scaffold, we reasoned that pioneer fascicles should exhibit molecular profiles that are distinct from surrounding tissue. We focused on cell adhesion molecules (CAMs) because they mediate contact-dependent axon–axon interactions and have been implicated in pioneer fascicle outgrowth and nerve ring patterning^20,34–38^, consistent with our observations of SAX-3/Robo enrichment at rosette centers (Fig. 3b-c). Using published embryonic scRNA-seq data^19^, we assessed the correlation of CAM expression across all annotated CAM genes — matching the gene set used by^20^ — among pioneers and surrounding tissue (largely corridor cells; Table S3, ‘CAM_similarity’). Pioneers show higher pairwise CAM correlation with one another than with corridor cells (mean r≈0.57 vs. 0.40, Fig. 5d), indicating pioneers are broadly molecularly distinguishable from surrounding tissue.

However, coordinating follower placement requires pioneers to be individually differentiable — not just distinct from surrounding tissue. Because the similarity measure is unable to resolve differences between pioneer pairs, we turned to a discordance measure restricted to the 50 most variable CAMs, since discordance among broadly and uniformly expressed genes is not informative. When pioneers are ordered by their eventual anterior–posterior position of their fascicles within the nerve ring, CAM discordance increases with pairwise-distance, revealing a graded pattern of molecular differentiation (Fig. 5e; median ∼3.8 differing CAMs per shared CAM across all pairs; Table S3, ‘embryo_cam_discordance_table’) that could support affinity differences among spatial domains.

### Follower axons track the pioneer scaffold to establish spatial domains

Both EM and live imaging demonstrate that several follower axons (AVA, AIY, RMDD among others) initiate outgrowth before pioneer fascicles have met at the dorsal midline. AVA and RMDD extend directly along the ProxL and SubL fascicles, respectively (Fig. S5f-h, Movie S5). AIY shows a more striking behavior: growth cones are projected in variable directions until the SMDD pioneer fascicle is encountered, at which point outgrowth proceeds directly along it (Fig. 5f, Movie S5). Together, these observations indicate that follower axons actively seek out the pioneer scaffold rather than simply growing alongside whatever is nearby.

Follower outgrowth relative to the scaffold is staggered and not fixed. Despite being among the first followers to reach the scaffold, AIY makes variable contacts with pioneers in larval EM reconstructions (Table S4, ‘aiy_contact’), consistent with later followers intercalating between pioneers and early followers as assembly proceeds (see Methods for EM contact-network definitions and thresholding limitations).

To assess how pioneer-follower interactions could give rise to nerve ring spatial domains, we asked whether followers sharing conserved contact with a common pioneer in the larval EM data are more likely to share domain membership than followers linked to different pioneers (a follower may share contact with more than one pioneer). Followers sharing a common pioneer contact share domain membership significantly more often than expected under a permuted null (observed: 0.51; null mean: 0.38, s.d. 0.046; Fig. 5g), while followers linked to different pioneers share domain membership significantly less often than expected (observed: 0.26; null mean: 0.31, s.d. 0.015; Fig. 5g). This probabilistic — rather than deterministic — relationship between pioneer contact and domain membership is consistent with the stochastic assembly process described throughout.

These observations suggest that pioneers serve as local organizers of follower outgrowth thereby constraining which followers make mutual contact (Fig. 1). We refer to this constraint as an axon’s neighborhood: the set of axons that contact a given axon when interactions are considered across a population of animals — not necessarily within any single individual, nor with every pairwise contact realized in every animal. In practice, this constraint is indiscernible within an individual, but is quantifiable across a population. We hypothesize that neighborhoods, shaped by early morphogenetic events, define the space within which pairwise interactions can occur, potentially reconciling stereotypy (repeated availability of contacts across individuals) and individuality (stochastic variation in which contacts occur). Whether constraining axon interactions to local neighborhoods explains the wiring statistics observed empirically is a quantitative question that can neither be resolved from static anatomy nor targeted perturbations (e.g. loss-of-function) alone. We address it directly next, with an agent-based model that quantifies how neighborhood constraints bias population-level wiring patterns.

### Modeling nerve ring innervation reveals constrained axon neighborhoods

We used agent-based simulations to test an organizing framework where individual contacts are stochastic outcomes within morphogenetically constrained neighborhoods. Two patterns are evident in larval EM reconstructions. (i) Axons organize into spatial domains where axons in the same domain make more contact than axons in different domains (Fig. 6a,b). (ii) Across a population, a given axon contact is most likely to occur in either one or all animals (Fig. 6c(i)) — a pattern that would be surprising if contact variability reflected unstructured developmental noise (Fig. 6c(ii)). Under our framework, these patterns emerge not from a molecular code but from the geometric logic of constraint: pioneer scaffolds, entry points and neighborhood boundaries that bias wiring without specifying individual contacts.

**Figure 6:**
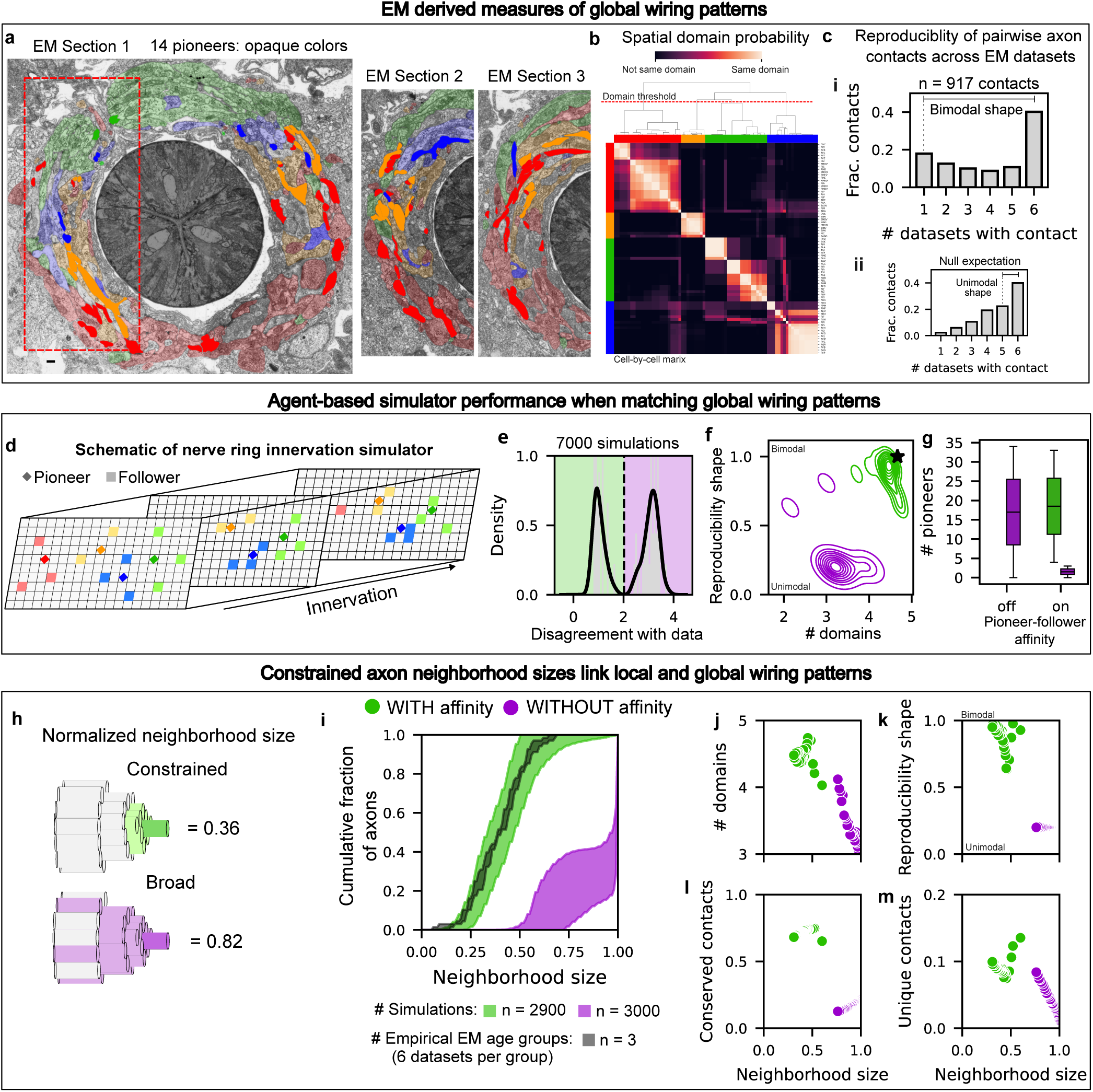
Constrained axon neighborhoods link local interactions to global nerve ring structure. **a**, Three representative EM sections from the L4 nerve ring along the posterior–anterior axis. Neurons colored by adult domain assignment; opaque coloring marks pioneers identified here. Left and right sides analyzed separately. Sections 2, 3 correspond to the dashed, more anterior region. Scale bar, 1 µm. **b**, Spatial domain organization of reproducible adult axon contacts. Matrix entries show the probability (0–1) that a pair of cells share a domain, via population-based clustering. Red dashed line: hierarchical clustering cut-off. Color bars denote domain membership. **c**, Reproducibility of pairwise axon contacts across six bilateral datasets. (i) Empirical distribution is bimodal, reflecting contacts conserved across all datasets or present in only one. (ii) Unimodal distribution expected if variability reflected only developmental noise. **d**, Conceptual schematic of the agent-based self-organizing model. Pioneer axons (diamonds) establish a scaffold; follower axons (squares) innervate locally as pioneers extend posterior to anterior. Local interactions follow simple affinity rules. The schematic illustrates model logic rather than exact simulation geometry. **e**, Distribution of simulation distance metric (SDM) values across local self-organizing simulations. SDM quantifies simulated-empirical disagreement in global wiring patterns. Dashed line: kernel density–estimated threshold separating low-(magenta) and high-agreement (green) simulations. **f**, Low- and high-SDM simulations plotted by number of spatial domains (x) and reproducibility shape (y). Black star: empirical mean. **g**, Simulations grouped by pioneer number and affinity (on/off). Low-disagreement simulations are green; high-disagreement simulations are magenta. **h**, Schematic of axon neighborhood sizes. Neighborhood size: number of distinct axons accessible to a given axon during innervation, normalized by total axons minus one. Constrained neighborhoods are green; broad neighborhoods are magenta. **i**, Cumulative distributions of neighborhood size for simulations (with or without affinity) and empirical data. Curves show the fraction of agents (simulated) or neurons (empirical) with neighborhood size a given value. Shaded regions: 1 SD age-group composite graphs (Methods). **j**–**m**, Relationships between neighborhood size and: **j**, spatial domains; **k**, reproducibility shape; **l**, conserved contacts; **m**, unique contacts. Green points: simulations with pioneer–follower affinity; magenta points: without. Each point: mean across 100 runs per parameter set (pioneer number, affinity on/off). See Fig. S9 and Methods for complete distributions.

To test whether this logic explains these patterns, our agent-based model mimics nerve ring innervation (Fig. 6d): non-branching, cable-like axons innervate within a fixed volume, enter at empirically constrained locations with defined orientation (set by assumed guidance), and interact locally according to simple affinity rules (Movies S6, S7). In principle, either constraint or specificity alone can explain why some contacts are highly conserved. We therefore evaluated the model against the full reproducibility distribution — spatial domain organization and bimodal reproducibility (Table S4) — rather than individual axon contacts. Although embryonic ultrastructural reconstructions are not yet available, these are age-invariant properties observed in larval connectomes (Fig. S6), providing stable benchmarks.

The simulator focuses on a local self-organization framework in which follower axons stochastically innervate around pioneers (Fig. S7; Table S5, ‘lso_basic_info’). To establish baseline performance, we varied the number of pioneers, gave each follower affinity to at most one pioneer, toggled ‘on’ or ‘off’, while holding total axon number, entry points, and growth volume fixed. We quantified agreement between simulations and empirical benchmarks using a simulation distance metric (SDM).

Across parameter space, simulations segregated into regimes that either reproduced or failed to reproduce empirical nerve ring organization (Fig. 6e). Local self-organization robustly recapitulated both spatial domain structure and bimodal reproducibility (Fig. 6f) when pioneer-follower affinity is ‘on’ and the tissue has a minimal pioneers density (>5; Fig. 6g), consistent with the 14 rosette-derived pioneers observed in vivo.

Successful simulations were characterized by constricted axon neighborhoods (Fig. 6h,i), the intermediate link anticipated above. Constricted neighborhoods coincide with limited spatial domains (Fig. 6j), bimodal reproducibility (Fig. 6k) and increased fraction of conserved (Fig. 6l) or unique (Fig. 6m) pairwise contacts (Fig. S9a-d). Together, these effects indicate that limiting neighborhood size facilitates precision while still permitting individuality. Neighborhood sizes measured from EM matched simulations with constricted neighborhoods (Fig. 6h), supporting a causal role in shaping wiring patterns.

An alternative positional-specification model — in which axons respond to fixed 3D coordinates rather than local pioneer affinity — only reproduced nerve ring organization under implausibly low response variability (Fig. S7b; Table S5, ‘pi_basic_info’), reinforcing that local interaction, not predetermined position, governs neuropil organization (see Methods).

Together, these findings identify distributed pioneers and local axon–axon affinity as a parsimonious framework — one that does not exclude alternative molecular affinity mechanisms — for the coexistence of stereotyped and individualized wiring, providing a quantitative bridge between developmental morphogenesis, constrained axon entry, and emerging axon neighborhoods.

### Spatial and molecular constraints robustly limit axon neighborhood size

We next asked which local interaction constraints are required to robustly limit pioneer-follower neighborhoods during nerve ring assembly (Fig. 7a). Guided by the rosette-based entry architecture, we added three parameters to the simulator: (i) follower affinity specificity (Fig. 7b), (ii) pioneer-follower proximity at nerve ring entry (Fig. 7c), and (iii) molecular distinctiveness among neighboring pioneers (Fig. 7d, Table S5, ‘lso_robust_info’). Together, follower specificity and pioneer distinctiveness constitute the model’s molecular specificity. We define robustness as insensitivity to parameter perturbation and therefore tolerance to developmental stochasticity. Modest changes in interaction rules should not shift agreement with empirical benchmarks from good to poor.

**Figure 7:**
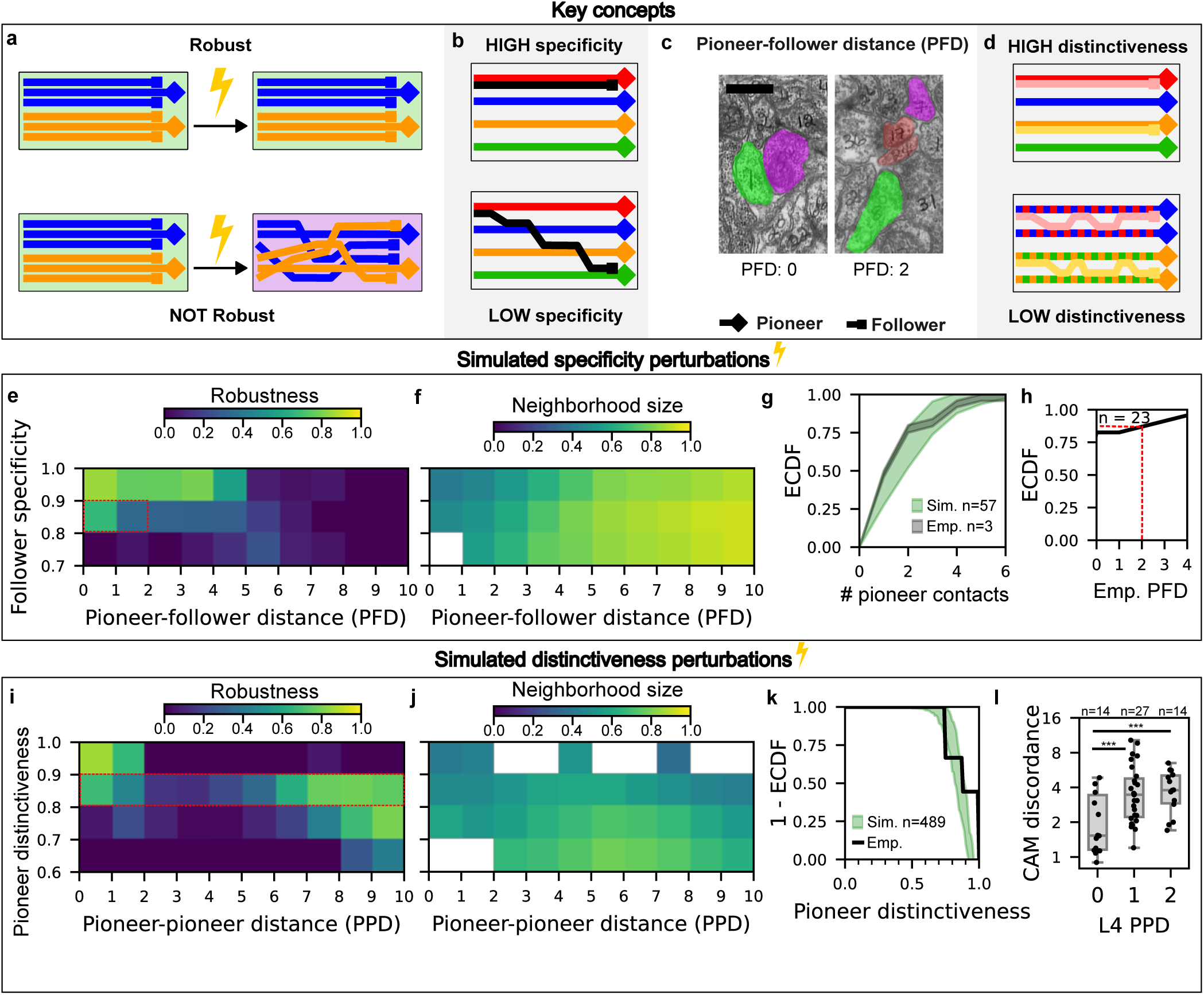
Spatial and molecular constraints define robust axon neighborhoods. **a**, Robustness assay: simulations matching empirical data (low SDM, green) are perturbed in parameter space and repeated. Robust simulations remain low SDM; non-robust simulations shift to high SDM (magenta). **b**, Follower specificity: number of pioneers attracting a given follower. **c**, Pioneer-follower distance (PFD) in L4 EM: number of axons separating a pioneer (magenta) from a follower (green) at nerve ring entry (0 = adjacent, 2 = two intervening axons). **d**, Pioneer distinctiveness: how distinguishable pioneers are to followers; low distinctiveness permits attraction to multiple pioneers. **e**, Robustness heatmap: simulations varying follower specificity (y) and PFD (x). Robustness is the fraction of 999 perturbation runs retaining low SDM. **f**, Corresponding heatmap of average neighborhood size across perturbed-follower simulations. **g**, Cumulative distributions of follower specificity for simulations (green; *n* = 57 runs from parameter regime marked by red box in **e**) and empirical data (black; *n* = 3 reduced composite age-group datasets). Shading indicates ± SD. **h**, Cumulative distribution of PFD in the empirical nerve ring; red dashed line: fraction of followers with PFD ≤ 2 (*n*: number of followers in reduced empirical network). **i**, Robustness heatmap: pioneer distinctiveness (y) and pioneer-pioneer distance (PPD; x), 999 runs. PPD defined analogously to PFD. **j**, Corresponding heatmap of average neighborhood size across perturbed-pioneer simulations. White boxes (**f**,**j**): regimes insufficiently sampled to estimate neighborhood size. **k**, Reverse cumulative distributions of pioneer distinctiveness: simulations (green; *n* = 487 runs, regime marked by red box in **i**) and empirical data (black; single composite distribution, since CAM data provide one average expression profile per pioneer, not replicates; Methods). Shading indicates ± SD. **l**, Pairwise CAM discordance between pioneers as a function of minimal separating axons (PPD). Boxes show median and interquartile range; dots indicate individual pairs. Welch’s *t*;-test, *p* < 0.01.

Assembly was most robust when followers, on average, exhibited affinity for fewer than ∼20% of pioneers and entered the nerve ring within ∼4 axon diameters of its matched pioneer, yielding constricted neighborhoods (Fig. 7e,f). Broader affinity or more distant entry produced high SDM scores, failing to reproduce spatial domains and bimodal reproducibility. EM analysis confirms both constraints. Because follower specificity cannot be measured directly *in vivo*, we estimated it using above-average EM contacts with the 14 pioneers (Fig. S8a), yielding a reduced contact network (Fig. S8b). Within this network, most followers contacted only 1–4 pioneers (Fig. 7g, Fig. S8b, Table S6, ‘follower_specificity’), consistent with the model’s predicted specificity regime, which also predicts followers should enter proximal to their matching pioneer. EM data confirms that ∼78% of followers entered the nerve ring within two axon diameters of their pioneer (Fig. 7h; Table S6, ‘pioneer_follower_distance’), identifying entry position as a key control point.

We next asked how molecular distinctiveness modulates neighborhood size under dense pioneer packing. In the model, molecular distinctiveness is represented as a generic differentiability among pioneers — not tied to specific molecular cues — that determines how easily followers discriminate between them. Simulations uncovered a trade-off between physical spacing and molecular distinctiveness (Fig. 7i). Indistinguishable neighboring pioneers required spatial separation, whereas sufficiently distinct pioneers could be placed adjacently, since molecular contrast alone preserves constricted neighborhoods (Fig. 7j). This trade-off predicts that molecular differences among pioneers should be organized at a scale sufficient to preserve neighborhood separation *in vivo*.

To test whether such molecular distinctiveness exists *in vivo*, we used CAM discordance metrics derived from embryonic scRNA-seq data^19^ to quantify differences among pioneers (Fig. 5e). We define molecular distinctiveness as the fraction of pioneers whose CAM discordance is below a minimal threshold (Methods). Pioneers are typically molecularly similar to at most one other pioneer, yielding intermediate distinctiveness scores (0.8–0.9; Fig. 7k, Fig. S8d), consistent with the model’s prediction that spatially proximal pioneers should remain distinguishable (Fig. 7i). Integrating L4 scRNA-seq^20^ with EM reconstructions^13^ showed proximal pioneers differ by ∼3 CAMs per shared CAM, versus ∼8 for distal pairs (Fig. 7l, Fig. S8g; Table S3, ‘l4_cam_discordance_table’). Together with our embryonic data (Fig. 5e), these results indicate that graded differentiation supports constrained neighborhoods.

Together, early morphogenesis and graded differentiation define the spatial and molecular conditions under which pioneer-follower self-organization reliably reproduces nerve ring architecture across individuals.

### Moderate interaction specificity within constrained neighborhoods expands reproducible wiring

A central question remains: how can neighborhoods stay constricted while followers associate with more than one pioneer? Intuitively, broader affinity should expand neighborhoods and undermine reproducible organization.

Simulations show that this intuition holds only beyond a narrow regime. Moderate specificity — where followers are attracted to ∼10–20% of pioneers — drives neighborhood sizes to the upper bound tolerated by robust organization, but does not exceed it (Fig. 8a). Neighborhood sizes in this regime overlap with those from high-specificity simulations (single-pioneer affinity) and with the empirically derived neighborhood size distribution from a reduced EM network (Fig. 8a). Robust self-organization therefore tolerates flexible local interactions so long as effective neighborhoods remain constrained.

**Figure 8:**
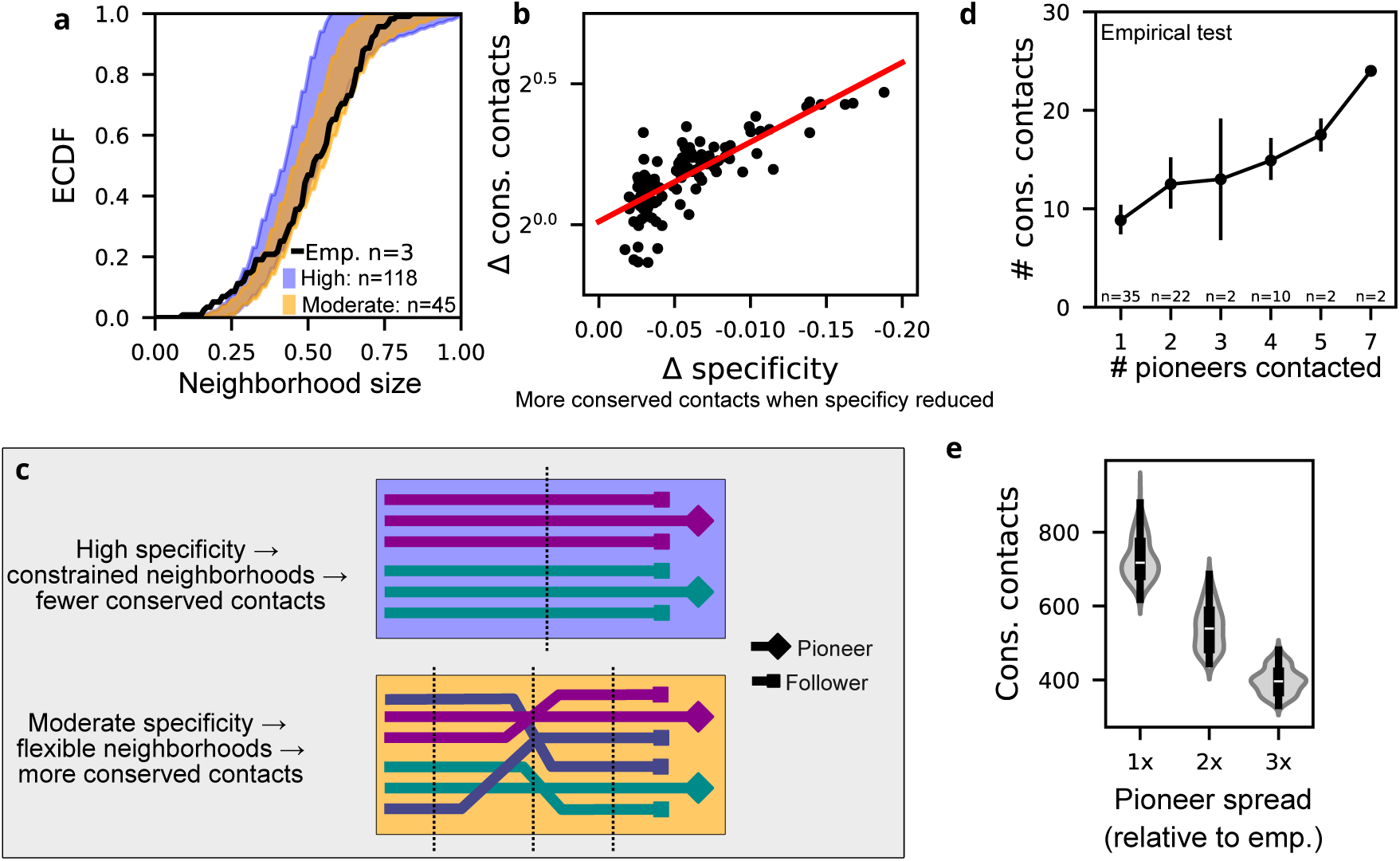
Flexibly constrained axon neighborhoods increase conserved wiring. **a**, Cumulative distributions of normalized neighborhood size for high-specificity (*n*=118) and moderate-specificity (*n*=45) simulations and for the reduced empirical network (*n*=3 age groups). Shading indicates ±1 SD for simulations; empirical curve shows the mean only. Neighborhood size is normalized by total neuron number. **b**, log fold change in the fraction of conserved contacts when follower specificity is reduced from high to moderate. Each point compares matched simulations with identical initial conditions (*n*=108 paired comparisons). Red line, linear fit. **c**, Conceptual schematic illustrating how reduced follower specificity expands neighborhood overlap, increasing the probability of reproducibly conserved contacts across individuals. Top: high-specificity fascicle, growing along a single, consistent path. Bottom: lower-specificity fascicle; dashed lines mark points where innervation transitions expand the axon neighborhoods. **d**, Empirical test of model prediction. Conserved contacts as a function of the number of pioneers contacted (proxy for follower specificity). Points show mean ± SD. Sample sizes (neurons): 1–5,7 pioneers contacted; *n*=35, 22, 2, 10, 2, 2, aggregated across age groups. Sample sizes decrease at higher values because fewer neurons contact many pioneers (Fig. 7g). **e**, Increasing pioneer spatial spread reduces conserved contacts. Violin plots show conserved-contact distributions as a function of relative pioneer spread in the transverse plane. Pioneer spread is quantified as simulated aspect ratio divided by the empirical aspect ratio along the transverse plane (e.g., 2× indicates twice the empirical pioneer spread). White dot, median; gray bar, interquartile range.

Within this constrained neighborhood regime, reducing specificity led to a logarithmic increase in population-conserved contacts (Fig. 8b,c). This increase arises from additional contacts among pioneer–follower pairs and shared-pioneer followers, alongside a reduction in contacts between followers linked to different pioneers (Fig. S9e-g). This redistribution matches the well-defined spatial domains observed empirically (Fig. 5d, Fig. 6b): favoring interactions among axons sharing a pioneer concentrates contacts within restricted regions while suppressing interactions across pioneer-defined boundaries. While systematic counterfactual manipulation of wiring specificity *in vivo* is not currently possible, followers contacting more pioneers — a proxy for lower specificity — do show more conserved contacts (Fig. 8d), consistent with the model’s prediction. Moreover, followers sharing a pioneer show greater conserved contact frequency than followers linked to different pioneers (*p*=6.73e-4, Fig. S9h), consistent with pioneer-defined neighborhoods as a primary organizer of contact reproducibility across individuals.

However, relaxing geometric confinement — spacing pioneers further apart than observed *in vivo* — reduced conserved contacts and degraded robustness even at high follower specificity (Fig. 8e, Table S5, ‘lso_spread_info’). Together, these results indicate that moderate specificity enhances stereotypy and domain organization only when axon growth is confined to a sufficiently restricted volume, as enforced by corridor-cell-restricted pioneer trajectories (Fig. 4), analogous to the spatial constraint modeled here.

Together, these results identify constrained neighborhoods as the mechanism reconciling flexible local interactions with robust, reproducible wiring — limiting the effective interaction space while permitting multiple local associations, and linking morphogenetic scaffolds to emergent network organization.

## Discussion

Biological development is a stochastic process that produces varying phenotypes^39^. Neural wiring is no exception^40^, but connectomic studies are revealing that wiring variability can be structured as a mix of highly reproducible and individualized connections across a population^13^. Hence, wiring reproducibility is a spectrum, where different species can arrive at their own mix of highand lowreproducible contacts. By accounting for this spectrum, rather than a single representative pattern, we arrive at a richer developmental account of wiring assembly — where neural wiring can leverage a constraint-specificity trade-off to achieve a diverse range of reproducibility.

Our empirical observations fill in key morphogenetic gaps in early *C. elegans* nerve ring development. Initial cell placement follows the stereotyped cell lineage^15,41^, subsequently refined through head involution and neural cord convergent extension^28,42^. Cells along the nerve ring path converge to form rosettes, partially mediated through SAX-3 (Fig. 3b-d), which prefigure entry points (Fig. 2a). Pioneer fascicles project from rosette centers, oriented by long-range cues from surrounding tissue^24–27,43^ but also constrained by the corridor cells linking subsequent rosettes (Fig. 4). The pioneer fascicles are molecularly distinct from surrounding tissue (Fig. 5d), self-sort into a scaffold that corresponds to domain organization (Fig. 5c), and locally organize follower innervation (Fig. 5f,g). This physical scaffold is a broadly predictive prerequisite of synaptic partnership^44^, yet it does not fully specify the number^45^ or subcellular placement^46^ of synapses. Whether these synaptic decisions are constrained by the early scaffold or specified independently by later mechanisms remains an open question.

Rosettes, corridor cells, and the pioneer scaffold constrain later wiring choices, but we needed to integrate them into a computational model to explain population wiring reproducibility. Our agent-based model implements the same three constraints: (i) restricted entry (rosettes), (ii) constrained trajectories (corridor cells), and (iii) local self-organization of followers around pioneers. However, our model necessarily simplifies biological complexity. Guidance and patterning in the neuropil are treated as separable mechanisms, with long-range axon guidance taken as a given. Axons are modeled as simple cables in 3D space and do not consider the more elaborate morphologies of some axons^13,17^, which likely results in an overestimate of empirical neighborhood sizes. Nevertheless, by comparing our model to the nerve rings, we elevate these empirical findings to a falsifiable prediction: morphogenetic constraints and molecular specificity trade off to organize robust yet flexible wiring.

Our model identifies the physical regimes where this trade-off robustly occurs. Morphogenetic constraints (spatial confinement, entry precision) set an upper bound on neighborhood size — exceeding it breaks robustness (Fig. 7f,j). Within that bound, spatially proximal pioneers need to be sufficiently differentiable by followers (∼3 CAMs per shared CAM for close pioneers vs. ∼8 for distant ones, Fig. 7i-l) and lower follower specificity is tolerated unless it yields larger neighborhoods (followers can have affinity for 1-4 pioneers; Fig. 7e-h). Counterintuitively, lowered specificity benefits the neural circuit: restricting the space where axons can innervate expands the repertoire of conserved contacts across a population (Fig. 8a-c). Our findings do not require a specific molecular implementation of this constraint — they define the conditions any such system must satisfy. Cell adhesion molecules are a plausible implementing class^47^: differential CAM expression among pioneers maintains neighborhood boundaries while permitting overlap (Fig. 7l), consistent with the graded molecular differentiation observed across the scaffold (Fig. 5e). SAX-3 illustrates both the promise and difficulty of this approach: its punctate enrichment at rosette vertices is more consistent with adhesion than classical guidance function (Fig. 3b,c), yet *sax-3* loss-of-function affects rosette cohesion and fascicle orientation only partially independently — underscoring that even well-studied CAMs resist clean mechanistic categorization from expression and localization data alone (Fig. S3); identifying these CAMs via loss-of-function remains important future work.

While our trade-off prediction was derived in a simple nervous system, it can potentially be generalized to more complex systems. Broadly, tightly constrained developmental systems should show bimodal contact reproducibility, while systems with relaxed constraints should show flatter, more uniform distributions. This logic yields a specific prediction for more elaborate neurons. Branches lack an obvious analog to the rosettes and corridor cells that constrain the nerve ring^40^, and in this loosely confined regime, our model predicts that even high molecular specificity cannot rescue reproducibility at the level of individual axon-to-axon contacts (Fig. 8e). This is consistent with the field’s general expectation that highly branched neurons connect stochastically as individuals, with stereotypy instead emerging at the level of neuron classes^48^. This highlights that the degree of wiring reproducibility can vary not only across species but also across topological scales within the same organism. A key takeaway may be that the constraint-specificity trade-off is the general relation that lets each developmental context find its place along the reproducibility spectrum.

## Methods

### *C. elegans* strains and genetics

*C. elegans* strains were raised at room temperature following the standard protocol^12^. Strains and genotypes used are listed in Table S1.

### Embryo Preparation for Imaging

*C. elegans* embryos were obtained by dissecting gravid adults in M9 buffer and staged at the 2– 4 cell stage. Embryos were mounted in M9 with 18.8 μm polystyrene beads, which prevented compression, and sealed between coverslips with petroleum jelly to avoid evaporation during long-term imaging^16^.

### Embryogenesis Imaging

Embryos were imaged on either (1) a Zeiss AxioObserver Z1 inverted microscope with Yokogawa CSU-X1 spinning disk, Olympus UPlanSApo 60× silicone oil objective, VisiTech laser merge module, and dual Hamamatsu C9100-13 EMCCDs, or (2) an Olympus inverted microscope with VisiTech iSIM and Olympus UPlanSApo 40× silicone oil objective. Images were acquired with Metamorph at an ambient temperature of ∼20 °C. For each embryo, 35 optical sections (0.75–1.0 μm spacing) spanning the whole embryo were collected every 1–3 min for up to 400 min. Typical datasets covered 250–400 min of development.

### Use of axon terminology

For simplicity, we refer to the primary processes entering the nerve ring as axons, acknowledging that *C. elegans* neurons do not exhibit the classical dendrite–axon polarity seen in vertebrates.

### Rosette and corridor cell identification

Rosette and corridor cells were identified by lineage tracing from the 2–4 cell stage using StarryNite to track nuclear divisions labeled by ubiquitously expressed histone-RFP, followed by manual correction in AceTree^49–51^. Cells were further characterized by morphology changes using a broadly expressed UNC-33::GFP membrane protein. Rosettes were defined as convergence of four or more cells to a central vertex. Rosette composition was corroborated using sparsely expressed membrane markers (LIM-4::GFP, CND-1::GFP). We identified both precursor and final rosettes (Fig. S1, Movie S1). Final rosettes form just prior to fascicle outgrowth (Fig. 2a, Movie S3); the nerve ring served as a fiducial point to trace pioneer axon outgrowths back to their rosettes. Precursor rosettes form transiently prior to the final rosettes. All rosette-based quantification in this study (SAX-3 enrichment and sax-3 phenotyping) was performed on final rosettes. We conjecture that precursor rosettes may aid local cell positioning and migration, but our data does not test this directly.

### SAX-3/Robo localization quantification

SAX-3::GFP fluorescence was quantified manually in FIJI^52^. For each rosette center or neighboring cell-cell junction, mean per-pixel intensity was measured within a manually drawn bounding oval across up to three consecutive z-slices centered on the structure of interest; where signal was confined to a single z-slice, that slice was used. Background intensity was estimated from a region outside the embryo and subtracted from all measurements. To account for day-to-day variation in imaging conditions, enrichment ratios were computed by dividing background-subtracted rosette center or junction intensity by the mean background-subtracted intensity of the surrounding cell bodies within the same image. Measurements were performed at SubL, ProxL, and DistL rosette centers and neighboring cell-cell junctions across three embryos (*n*=11 rosette centers, *n*=21 junctions). Enrichment ratios were averaged across z-slices for each measurement (Table S2, ‘SAX3_localization’). Differences between rosette centers and neighboring junctions were assessed using a two-tailed Welch’s t-test on pooled measurements across all three rosette types.

### *sax-3*(ky123) rosette and fascicle analysis

We crossed *sax-3*(ky123) with a strain containing an integrated UNC-33::GFP membrane marker (Table S1) and imaged 21 embryos. Head morphology was assessed independently of, and prior to, rosette and fascicle scoring. Embryos with abnormal head morphology (*n*=3) were excluded, as rosette and fascicle status could not be reliably interpreted in the context of broader head malformation. In the remaining embryos, image stacks were rescaled to isometric voxels. Rosette formation was assessed by cell vertex convergence at anticipated locations, based on comparison with wild-type UNC-33::GFP embryos of matched orientation. Wild-type embryos were imaged using an extrachromosomal marker (Table S1), which was brighter and easier to use for establishing baseline structures. Because the extrachromosomal and integrated markers differ in brightness, fascicle presence/absence calls were validated using internal controls within and across *sax-3* embryos independent of any comparison to wild type. For example, we would look for asymmetric presence or absence of the same structure on the left versus right side of a single embryo. Fascicle presence was scored by increased fluorescence intensity along the nerve ring path, consistent with multiple neurite membranes aggregating together during innervation. To confirm absence was not due to viewing angle, embryos deemed to have absent fascicles were further examined via orthogonal projections — image slices parallel to the direction of neurite growth. Decreased fluorescence intensity only tracks the behavior of the neurite fascicle as a whole, not individual neurites. We therefore cannot preclude the possibility that some individual neurites are correctly oriented even when aggregate fascicle signal is absent.

### Corridor cell ablation

We ablated the corridor cells that link the SubL and ProxL rosettes. Using automated cell tracking ablation software ShootingStar^33^, we simultaneously eliminated sister corridor cells IL2V and IL1Vm (the mother of IL1V) by identifying and laser ablating their mother cell: left/right cell homologs ABalppapp/ABarappp. As an internal control, we only ablated the mother cell on one side of the embryo and checked for normal development on the opposite side.

Fascicle outgrowths were monitored using membrane marker LIM-4::GFP, which is sparsely expressed in SubL and ProxL rosette cells (Table S2, ‘pioneer_markers’). Following ablation, SubL outgrowth was monitored by time-lapse imaging to characterize the resulting phenotype. In most successful ablations, SubL projections initially stalled before extending — most wandered off the trajectory seen in mock ablations, but one embryo recovered correct outgrowth (Fig. S4). To check for off-target effects, we checked for normal cell cycles of the non-ablated mother cell on the contralateral side (or similarly placed cell if the contralateral cell could not be easily tracked) and of 3-4 cells physically proximal to the target cells (Table S2, ‘ablation_validation’). Ablation of the corridor cell mother also eliminates cells BAG and SMDV (Fig. 4b). However, these cells are not corridor cells, are not located between the SubL and ProxL rosettes, and we do not observe that these cells directly engage in any of the rosettes of this study.

We used a binomial statistical test to determine if the ablation results were significant (Fig. 4e). The null hypothesis is that the fraction of animals with WT axon growth is not (statistically) distinguishable between mock ablated, proximal ablation and ablated animals. To determine if the different groups are distinguishable, we assume samples are drawn from a binomial distribution defined parameters *p* (the fraction of animals with WT axon growth) and *n* (the number of embryos). We define the mean as *np* and estimate the variance as *np*(1 − *p*) which can be used to compute the standard error. We then use the mean and standard error in a Student *t*;-test.

### Triangulating axon positions in the embryo

To determine the relative positions of axons within the early nerve ring, we triangulated axon positions across three worm strains, each expressing two of four membrane markers in different subsets of rosette cells (Fig. S5). The markers used were CEH-10 (RMED, ALA), ZAG-1 (AVD, AVJ), CND-1 (RIV, SAAV, SMDV, and unidentified cells), and LIM-4 (RIV, SAAV, SIAD, SIAV, SIBV, SMDD) (Table S2, ‘pioneer_markers’).

By comparing marker pairs, we inferred boundaries and relative positions of axon bundles. For example, CEH-10 and CND-1 delineated anterior-posterior boundaries corresponding to RMED and ALA, while CND-1 and ZAG-1 revealed medial positioning of AVD and AVJ. CND-1 fluorescence indicated two parallel axon bundles (anterior and posterior), and comparison with LIM-4 identified specific cells in the anterior bundle (SAAV, SIAD, SIAV, SIBV, SMDD). RIV, SAAV, and SMDV project from the ProxL rosette, allowing us to infer that RIV is in the anterior bundle.

To quantify relative axon positions, we used FIJI^52^ to measure distances between centers of axon bundles (Table S2, ‘pioneer_triangulation’). These measurements enabled triangulation of pioneer axon positions, which were then compared with EM reconstruction.

### Follower-pioneer time-lapse imaging

Follower outgrowth behavior relative to pioneer fascicles was imaged in AIY (TTX-3b::GFP) and RMDD (ZAG-1::GFP; CND-1::RFP), allowing direct observation of growth cone dynamics during outgrowth. No fluorescent marker was available for AVA at the time this dataset was collected, as EM data motivating its inclusion as a follower of interest was not yet available; AVA-pioneer contact was therefore assessed by EM reconstruction alone (see below, “Embryo reconstruction from electron microscopy”). RMDD, for which both fluorescence tracking and EM reconstruction were available, showed agreement between the two approaches: EM-confirmed pioneer contact corresponded to fluorescence-observed active tracking, and fluorescence-observed tracking corresponded to genuine ultrastructural contact rather than proximity between unmarked processes. This bidirectional agreement supports extending each single-modality inference to the evidence available for AIY (tracking behavior, fluorescence only) and AVA (genuine contact, EM only).

### Embryo reconstruction from electron microscopy

#### Electron microscopy of embryos

The EM data was generated by IK and published in collaboration with the ZB group^32^. We analyzed serialized EM datasets of 340 and 365 mpf embryos. Imaging resolution is provided in Table S1. Detailed protocols for embryo preparation and computational cell identification are described in the cited study.

#### Volumetric reconstruction

3d reconstruction from EM images was performed manually using TrakEM2^53^. Neurons with nerve ring outgrowths were prioritized, and peripheral cells were occasionally reconstructed to aid identification. Not all neurons are included. The resulting reconstruction datasets are available upon request.

### CAM profiling

#### Data source

CAM gene identities (141 total) and corresponding WormBase IDs were obtained from Table S3a of Taylor *et al.* (2021). Embryonic scRNA-seq data were downloaded from Packer *et al.* (2019). Following their published criteria, a gene was defined as “expressed” when the lower 95% confidence bound on the estimated transcripts was greater than zero. L4 scRNA-seq data were obtained from the CeNGEN consortium (“most permissive” expression matrix), with genes considered expressed when TPM > 0. Both datasets were used without modification. Analyses were restricted to CAM genes and identified pioneer neurons; for the embryonic dataset, only cells from the 390–510 mpf time window (follower innervation period) were included. Dataset links are listed in Table S3, ‘embryo_info’ and ‘l4_info’.

#### Dataset limitations

Both datasets resolve only neuron classes, not left/right homologs or dorsal/ventral subclasses of RME, SAA, SIA, SIB, and SMD. The embryonic dataset also lacks SAA cells within the 390–510 mpf window. Accordingly, we merged our resolved subclass identification in the composite contact graphs to match the single-cell dataset resolution.

#### Gene selection and binarized expression

We restricted our analysis to CAM-associated genes because these molecules are well-established mediators of selective axon–axon interactions^36^. Gene expression was binarized (1 = expressed, 0 = not expressed) to reduce sensitivity to scRNA-seq quantification noise and because quantitative CAM expression levels are difficult to interpret biologically within the context of cell-cell affinity.

#### CAM similarity between pioneers and surrounding tissue

Pairwise CAM similarity between cells was computed using the phi coefficient, a measure of association between binary variables equivalent to the Pearson correlation coefficient applied to binary data. For each pair of cells, CAM expression was represented as binary presence/absence vectors across all annotated CAM genes. The phi coefficient was calculated as

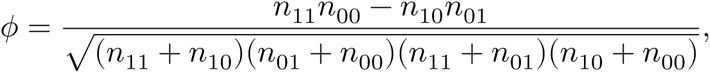

where *n*_11_ and *n*_00_ denote the number of CAMs jointly present or jointly absent in both cells, and *n*_10_ and *n*_01_ denote the number of CAMs present in only one of the two cells. This yields a symmetric similarity score ranging from −1 to 1, with self-similarity set to 1. Pairwise similarity was computed across all cells of interest (e.g., pioneers and corridor cells; Fig. 5d) to generate a full pairwise similarity matrix.

#### Pairwise CAM discordance

From the binarized matrix, we retained CAM genes expressed in fewer than 75% of pioneer neuron classes in each of the embryonic and L4 datasets (Fig. S8c), yielding 50 genes in each dataset. We reasoned that genes with intermediate expression frequencies carry the highest variance and thus the greatest discriminatory power. CAMs excluded by this process tend to be broadly expressed across multiple tissues and are primarily involved in general adhesion, guidance, or morphogenetic functions rather than selective wiring.

To quantify differences in CAM repertoires between neurons, we calculated a pairwise differential score:

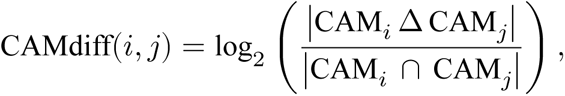

where Δ denotes the symmetric difference and ∩ the intersection of expressed CAMs between neurons *i* and *j*. CAMdiff scales by the differences in shared CAMs rather than total CAM counts. Unlike Jaccard similarity, this approach captures absolute differences between neurons, ensuring that neurons with similar relative overlap but differing numbers of CAMs are distinguished in a biologically meaningful way. Only binarized expression values were used, adhering to thresholds and definitions in the original datasets.

#### Empirical pioneer distinctiveness

We binarize the CAM pioneer by pioneer discordance matrix. Pairwise discordance below 0.3 is set to 1 and 0 otherwise, which converts the discordance matrix into a similarity matrix (Fig. S8d). For each pioneer, distinctiveness is defined as 1 minus the fraction of 1s in its row (excluding the main diagonal).

### Larval EM analysis

#### Dataset partitioning by developmental stage

To assess age invariance of the spatial domain and bimodal reproducibility benchmarks, we pooled nine larval EM datasets spanning L1 to adult (Table S4)^9,13^ and quantified benchmark stability across stages (Fig. S6d-f; Table S6, ‘sdm_metrics’). To reduce variance from developmental differences, datasets were grouped into three age categories: young (0, 5, 8 hr L1), middle (16 hr L1, L2, L3), and adult (L4 and two adults) (Fig. S6a, Table S4, ‘age_diff’). Each animal was further divided into left- and right-side axon sets, yielding six ipsilateral datasets per age group. Each dataset was distilled into a cell-by-cell contact matrix where matrix entry quantifies the amount of membrane surface area contact between axons of the two corresponding cells.

#### Contact graph generation

Contact graphs were derived from the contact matrices for each dataset, where nodes represent neurons and edges indicate pairwise axon contacts. As in previous analyses^13^, edges representing the smallest 35% of membrane contacts represent a small fraction of the total membrane contacts and are the least reproducible and therefore removed from the analysis.

#### Composite contact graphs

For each age group, the six ipsilateral graphs were merged into a composite contact graph. Each edge was assigned two attributes: (i) reproducibility frequency, the number of datasets in which the contact occurs, and (ii) conditional contact length, the average contact length measured only over datasets in which the contact exists. Summary statistics—including bimodality of contact reproducibility, number of spatial domains, and average axons per domain— were computed on these composite graphs (see below). Papillary neurons have trajectories that are orthogonal to the main nerve ring path^14^ and were excluded to maintain consistency with the agent-based simulations which only model the main nerve ring path. The resulting contact graphs are comprised of 68 neurons for all age groups and 817, 912 and 917 edges for young, medium and adult age groups, respectively (Table S4, ‘young_repro’, ‘middle_repro’, ‘adult_repro’)

#### Bimodal reproducibility shape

The reproducibility distribution is the histogram of reproducibility frequencies across all edges in the composite contact graph. Bimodal distributions of edge reproducibility (peaks at 1 and 6 datasets) are quantified as the reproducibility shape (v), defined as the normalized distance between the two highest peaks. Values near 1 indicate strong bimodality.

#### Spatial domains

Louvain community detection^54^ identifies spatial domains using edges with reproducibility frequency equal to 6 contact graphs, weighted by conditional contact length. Robustness to biological noise is addressed by repeating community detection 100 times with each iteration adding white noise to conditional contact length. We then determine consensus communities by creating a neuron-by-neuron matrix which quantifies how frequently neuron pairs were assigned to the same community. Final domains were then determined by performing hierarchical clustering on the neuron-by-neuron matrix (SciPy 1.0, fcluster) which generated a dendrogram. Neuron pairs within a dendrogram distance of 10 are assigned to the same spatial domain^13^.

The dendrogram distance threshold (10) was determined to be the coarsest threshold that produced consistent domain assignments across all 3 empirical age-group datasets and provides a principled basis of comparison between empirical and simulated data. Finer thresholds over-resolve the clustering into sub-domains whose boundaries are not reliably reproduced across individuals or simulation runs (e.g. compare young, medium and adult matrix intensities across Fig. 6b and S6b). Over-resolving reduces the reliability of the empirical-to-simulation benchmark (below) rather than sharpening it, because unstable sub-domain boundaries are not reproducible enough to serve as a reliable benchmark.

#### Neighborhood size

An axon’s neighborhood size is defined as the number of edges connected to it on the composite contact graph. Because edges can have varying reproducibility frequency, an axon’s neighborhood can include contact edges that only occur in a few datasets. Hence, neighborhood size characterizes the observed number of contacts across a population of animals. Operationally, to facilitate comparisons with simulations and with reduced empirical networks (see below) which have different numbers of axons, we normalize the neighborhood size by the total number of axons in the contact graph minus 1. Hence, the normalized version of the neighborhood size is the fraction of possible other axons that an axon could contact.

#### Approximating pioneer-follower affinity relations (“the reduced empirical network”)

Because we do not know the exact pioneer-follower affinity relations in the empirical nerve ring, we inferred pioneer-follower affinity based on above average contacts sizes. For each age group, we compute the distribution of conditional contact length between followers and pioneers (Fig. S8a). The reduced empirical network was then constructed from all pioneers and any other neuron that makes above average conditional contact with a pioneers, which were labeled as ‘followers’. Using this contact threshold, we approximated pioneer-follower assignments, where followers could be assigned to multiple pioneers (Fig. S8b). The resulting contact graph of the reduced network includes 14 pioneers and 23 followers, but we retain all of the associated contact edges between these axons regardless of whether or not they were above the conditional contact threshold.

#### Reduced empirical network thresholding limitations

Because later-innervating followers can intercalate between pioneers and early followers as assembly proceeds, some functionally meaningful contacts present during active outgrowth may fall below the above-average contact threshold used to construct the reduced empirical network, diluted by subsequent intercalation. This is a limitation of static contact thresholding rather than evidence against active tracking; live imaging and EM reconstruction are therefore complementary, with the former capturing transient growth-cone behavior during assembly and the latter offering contact resolution only after intercalation has already reshaped the network.

#### Pioneer-follower contact and domain membership

To assess whether followers sharing conserved contact with a common pioneer in the larval EM network are more likely to share nerve ring domain membership than followers linked to different pioneers, we used the reduced empirical network and pioneer-follower assignments described above. A follower could be assigned to more than one pioneer. For every pair of followers, we determined (i) whether the pair shared assignment to at least one common pioneer, and (ii) whether the pair shared nerve ring domain membership in at least one of the three age-group domain assignments (young, middle, adult), using the domain assignments described below (“Spatial domains”). Because domain assignment is derived from hierarchical clustering of the full axon contact network rather than from pioneer-follower contact structure, shared-pioneer-assignment status and domain-membership status are not circularly related.

We computed the observed fraction of same-domain pairs separately among follower pairs sharing a common pioneer and among follower pairs assigned to different pioneers. To generate a null distribution for each fraction, pioneer assignments were randomly permuted among followers 1,000 times, preserving the number of pioneer assignments per follower, and the same fractions were recomputed for each permutation. Observed fractions were compared to the resulting null distributions (Fig. 5g).

#### Pioneer-follower distance (PFD)

For each follower in the reduced network, the number of axons separating it from its nearest associated pioneer was manually assessed at the approximate point of nerve ring entry using L4 EM data. Rare cases where pioneer contact occurs later in the trajectory were labeled as separation >10 axons (Table S5, ‘pioneer_follower_distance’).

#### Pioneer-pioneer distance (PPD)

We define pioneer-pioneer distance as the number of edges separating pioneer nodes on the contact graph for the reduced network.

### Agent Based Simulator

#### Overview

The simulator model is intentionally minimal and does not attempt to capture the full biological complexity of nerve ring development. We assume long-range guidance gradients while omitting activity-dependent refinement, synapse formation, contralateral projections, and late-stage growth dynamics, focusing instead on early axon entry, local interactions, and geometric constraints revealed by embryonic imaging. This abstraction allows us to isolate which features are required to translate early morphogenesis into population-level wiring structure. As such, model success identifies a parsimonious class of mechanisms consistent with available data, while leaving open the possibility that additional biological processes refine or stabilize the pattern *in vivo*.

The agent-based simulator (ABS) models axon outgrowth in a simplified hexahedral grid representing one ipsilateral half of the nerve ring (Fig. S7a). The grid dimensions are 20 units width × 10 units height × 240 units length, matching empirical aspect ratio (Table S6, ‘aspect_ratio’). Each simulation contains 70 axons (corresponding to non-papillary head neuron classes), modeled as independent cable-like agents. At each step, axons advance one unit longitudinally and cannot move backward (Movie S6,S7).

Unlike previous models^45^, which used highly abstracted rules, our ABS incorporates biological realism through pioneer–follower dynamics and local spatial constraints, ensuring developmental fidelity while remaining computationally tractable.

#### Axon growth rules

Growth is asynchronous: at each step, axons are randomly ordered to prevent systematic bias. Each axon attempts to move into one of nine adjacent positions in the next slice. If the chosen site is occupied, it selects a neighboring empty site if available. If no empty site exists, it displaces the resident axon, with shifts propagating locally. Simulations run for 240 steps, with analysis restricted to the central 200 steps to minimize edge effects.

#### Contact Graphs, reproducibility and spatial domains

As with EM data, simulated axon interactions are quantified with contact graphs. Nodes = axons; edges = adjacency events: two axons occupy adjacent grid positions; edge weights = cumulative adjacency across simulation steps. Edges in the lowest 35th percentile are removed to mimic EM thresholding. Six replicate simulations per initial configuration are merged into a composite contact graph, matching the empirical dataset. Reproducibility shape and spatial domains are computed identically to EM analysis above (Larval EM analysis).

#### Simulation distance metric (SDM)

To compare simulated and empirical composite graphs, we defined a Simulation Distance Metric (SDM) based on three graph-level statistics: number of spatial domains (*d*), average axons per domain (*a*), and reproducibility shape (*v*). For each composite graph:

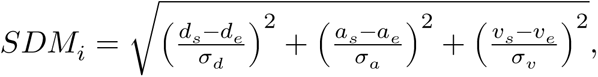

where *s* and *e* denote simulated and empirical values, and σ is the across-simulation standard deviation for each statistic. Normalization ensures equal weighting of *d*, *a*, and *v*.

#### Initial axon placement

To mimic corridor-cell restriction of scaffold pioneers, pioneer placement was restricted to a central subvolume of the grid at the first step. Followers were then placed adjacent to pioneers within the central subvolume, with any remaining followers positioned in surrounding regions. Each condition was repeated multiple times with randomized placements to marginalize initial-configuration effects.

#### Simulations models Baseline control: Positional specification model

Pioneers innervate first along fixed trajectories, establishing an initial scaffold. Follower axons then extend via a biased random walk guided by positional information. Following the classical conception of positional information^55^, we treat the morphogen gradient and the axon’s interpretation of that gradient as distinct. We assume a global positional signal uniquely specifies 3d spatial coordinates within the simulation grid. Rather than modeling the gradient explicitly, each follower samples a target position from a Gaussian distribution centered on its initial location, with standard deviation σ controlling positional uncertainty. Smaller σ corresponds to more reliable positional information and stronger bias toward the target, whereas larger σ introduces noise, increasing the likelihood of deviation from the intended trajectory (Fig. S7b; Table S5, ‘pi_basic_info’).

We tested up to 35 pioneers, each with a fixed trajectory, to evaluate how gradient fidelity and pioneer number influence follower placement and overall pattern reproducibility. PI models reproduce gross nerve ring geometry only under unrealistically tight anchoring, which is physically implausible given growth cone sensing limits^56^, the rapid timescale of follower innervation, and the limited physical space available within the neuropil to support steep, stable gradients. Across biologically plausible parameter ranges, PI models fail to capture the observed distributions of axon contacts (Fig. S7b).

#### Main model: Local self-organizing model via pioneer–follower affinity

In the baseline version of this model, pioneers extend first along fixed trajectories, and followers innervate via an unbiased random walk with no affinity toward pioneers (Fig. S7a,c-e; Table S5, ‘lso_basic_info’). This baseline isolates the effect of local interactions and provides a null control for reproducibility. Up to 35 pioneers were tested, each following a predefined trajectory within the 3d grid.

To incorporate pioneer–follower affinity, each pioneer and follower is assigned a binary (affinity) vector of length m, where m equals the number of pioneers. Pioneers are indexed and have a 1 at their index position and 0 otherwise. Followers are statistically assigned 1’s based on simulation parameters (see below). Each pioneer produces a local attractive field, modeled as a Gaussian (σ = 1 grid unit, maximum strength = 1) centered on its trajectory (Fig. S7a). At each step, a follower evaluates neighboring positions within the grid; the probability of moving toward a pioneer is proportional to the product of the attractive field and the binary overlap (shared 1s) between the follower’s and pioneer’s affinity vectors. When multiple pioneers influence a follower simultaneously, probabilities are normalized to sum to 1. Binary vectors are fixed for the duration of each simulation, and multiple 1s are allowed, reflecting partial affinity for multiple pioneers while maintaining local constraints. See Table S5 for parameter details and the publicly available simulation code for exact update and normalization rules.

This formulation isolates the contribution of local attraction while maintaining biologically interpretable variability in follower–pioneer interactions.

#### Varying follower specificity and pioneer–follower distance

Follower specificity was defined as 1 − (fraction of 1s in its affinity vector). High-specificity followers (low fraction of 1s) recognized few pioneers, often only one. Each follower was initially assigned to, on average, the th nearest pioneer, controlling pioneer–follower starting distance (Fig. S7f). When multiple affinities were assigned, additional pioneers were drawn from the neighborhood of the th nearest, keeping affinities spatially constrained. In general, specificity and pioneer distances are not uniform across agents, but randomly drawn from a normal distribution centered around a specified mean. This more naturally captures biological variance. See Table S5, ‘lso_robust_info’.

#### Varying pioneer distinctiveness and pioneer–pioneer distance

Pioneer distinctiveness was defined as 1 − (fraction of 1s in its affinity vector). Distinct pioneers contained only a single 1, corresponding to exclusive recognition. Non-distinct pioneers were generated by flipping 0s to 1s, introducing shared affinities among nearby pioneers. To control spacing between similar pioneers, flips were restricted to the d nearest neighbors (Fig. S7g). Similar to followers, distinctiveness and distances are drawn from distributions. See Table S5, ‘lso_robust_info’.

#### Robustness tests

To test minimal feature requirements for nerve ring organization, we simulated baseline conditions (≥5 pioneers, one-to-one pioneer–follower assignment, unique pioneer codes, nearest-neighbor placement). We then perturbed follower parameters (specificity, pioneer– follower distance) and pioneer parameters (distinctiveness, pioneer–pioneer distance), reran simulations with identical initial placements, and recalculated SDM. Perturbations maintaining SDM < 2 (Fig. 6e) were considered robust. In total, 3,000 follower-perturbation and 2,000 pioneer-perturbation simulations were performed (Table S5, ‘lso_robust_info’).

#### Contact increase with lowered specificity

We first simulate followers with affinity to only one pioneer and then repeat the simulation with lowered follower specificity. If both simulations are in the low SDM regime, we compute the change in follower specificity and number of conserved contacts (Table S5, ‘lso_robust_spec_change’).

#### Lateral pioneer spread model

Simulations are run while incrementally expanding the width of the cross-sectional area to allow pioneers to laterally spread further apart (Table S5, ‘lso_spread_info’). To maintain roughly the same cross-sectional area across simulations, we also reduce the height of the cross-section. We tested following cross-sectional areas (height x width): the empirical aspect ratio (10x20), 2x the empirical aspect ratio (7x28) and 4x the empirical aspect ratio (5x40).

### Parameter sampling and replication strategy

For each experiment, we sampled a large set of parameter combinations across predefined ranges (Fig. S7h, Table S5). Each parameter combination was evaluated using multiple independent stochastic realizations to account for variability arising from both initial conditions and growth dynamics. Specifically, simulations were initialized from multiple distinct random agent placements, and for each placement, six simulations were run with independent random seeds. This hierarchical replication strategy ensured that model outcomes were not driven by particular initial configurations or single stochastic trajectories.

### Quantification and statistical analysis

Details of quantification and statistical testing, sample size, center and dispersion are found in the figure legends and Method details section for individual analyses.

#### Aggregating simulation results across replicates

For Fig. 6d-e, distributions summarize SDM values across 7,000 simulations. Each of 70 parameter sets (varying 1–35 pioneers and pioneer– follower affinity on/off) was simulated 100 times; the mean SDM per set was calculated, and the distribution of these 70 means is shown.

#### Cumulative distribution function (CDF) analysis

To compare simulated and empirical distributions (neighborhood sizes, follower specificity, and pioneer distinctiveness), we analyzed their cumulative distribution functions (CDFs), which capture the full range and shape of stochastic variability and provide a more comprehensive assessment than tests based solely on means or variances (e.g., *t*-tests). For each measurement, a CDF was computed by calculating, for each value, the fraction of observations less than or equal to that value. To quantify variability across replicates, CDFs were computed separately for each replicate, and at each x-axis value the mean and standard deviation across replicates were calculated. Shaded regions in figures indicate ±1 SD across replicates, facilitating direct comparison of distributions across conditions or datasets.

## Author contributions

Conceptualization: CB, AS, WAM, HS, DCR, ZB. Project initiation and early experimental framework: ZB. Experimental design and data analysis: CB, KB, AS, ZB. Biological and imaging experiments: CB, KB, RC, LF, MWM. Reagent creation: MWM. Embryo EM dataset generation (imaging and collection): IK. EM reconstruction and analysis: CB, AS. Computational modeling and simulation analysis: CB. Research supervision: WAM, HS, DCR, ZB. Writing—original draft: CB. Writing—review and editing: CB, AS, with input from all authors. Corresponding authors: CB and AS

## Acknowledgments

We thank Lisbelle Adorno for administrative support. We thank members of the UNIL EM facility for technical assistance and the Caenorhabditis Genetic Center (funded by NIH Office of Research Infrastructure Programs P40 OD010440) for *C. elegans* strains. Research in the ZB, DCR, and WAM labs was supported by NIH grant R24-OD016474, and research in the HS lab was supported by the intramural research program of NIBIB, NIH and the Howard Hughes Medical Institute. Research in the ZB lab was further supported by an NIH center grant to MSKCC (P30CA008748). CB and AS were further supported by NIH grant R01GM152927. Research in the DCR lab was further supported by NIH grants R01NS076558 and DP1NS111778 and by an HHMI Scholar Award. HS and DCR acknowledge the Whitman and Fellows programs at MBL. AS was supported by grant 2019-198110 (5022) from the Chan Zuckerberg Initiative and the Silicon Valley Community Foundation. MWM was supported by NIH F32-NS098616.

## Resource Availability

### Lead Contacts

Correspondence and requests for materials should be addressed to Christopher Brittin or Anthony Santella.

### Material Availability

The strains generated in this study are available at the *Caenorhabditis* Genetics Center or by request from the lead contacts.

### Data and Code Availability

- Raw and processed imaging data and agent-based simulation outputs are deposited at Zenodo: https://zenodo.org/records/18704853
- Analysis code is available at GitHub: https://github.com/cabrittin/constrained-innervation-abm/
- External datasets used in this study are listed in Table S1.
- Movies S1–S7 and Supplementary Table Sx are not included with this preprint and will be available with the peer-reviewed publication.

---

**Figure S1:**
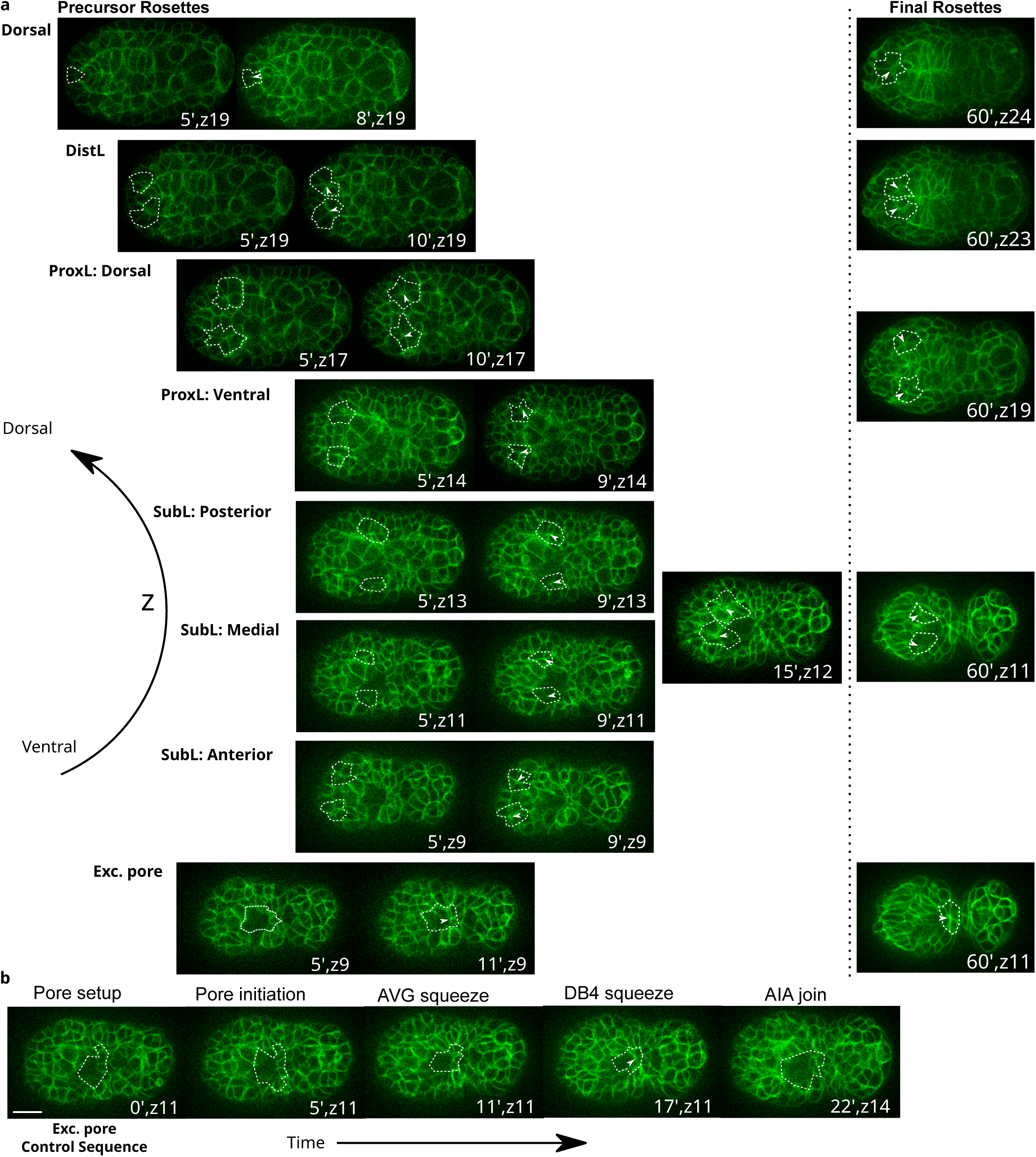
Spatiotemporal coordination of transient rosette formation during nerve ring assembly. **a**, Spatiotemporal map of the 8 nerve ring rosettes. Time proceeds left to right, synchronized to excretory (Exc.) pore development (350 mpf, see **b**) to establish a reproducible temporal landmark. Rosettes are ordered along the vertical axis (z) from ventral to dorsal slices of the embryo. Left of the dotted vertical line, sequential images show precursor rosettes before and after convergence; right of the line, a single image shows the corresponding final rosette prior to neurite outgrowth. White dashed line: outline of rosette cells. White arrowhead: rosette center. The SubL rosette first forms three intermediate rosettes (anterior, medial, posterior) that coalesce into a single rosette at minute 15. The ProxL rosette has two layers (ventral and dorsal) that converge around a central vertex. See Movie S1 for further details. All images are from the same ventral-facing embryo, anterior to the left. Scale bar: 10 μm. Slice thickness: 1 μm. **b**, Ventral portion of the Exc. pore rosette undergoes a stereotyped sequence of cell movements, used as an internal reference for rosette timing. See Fig. S2a,b for further details.

**Figure S2:**
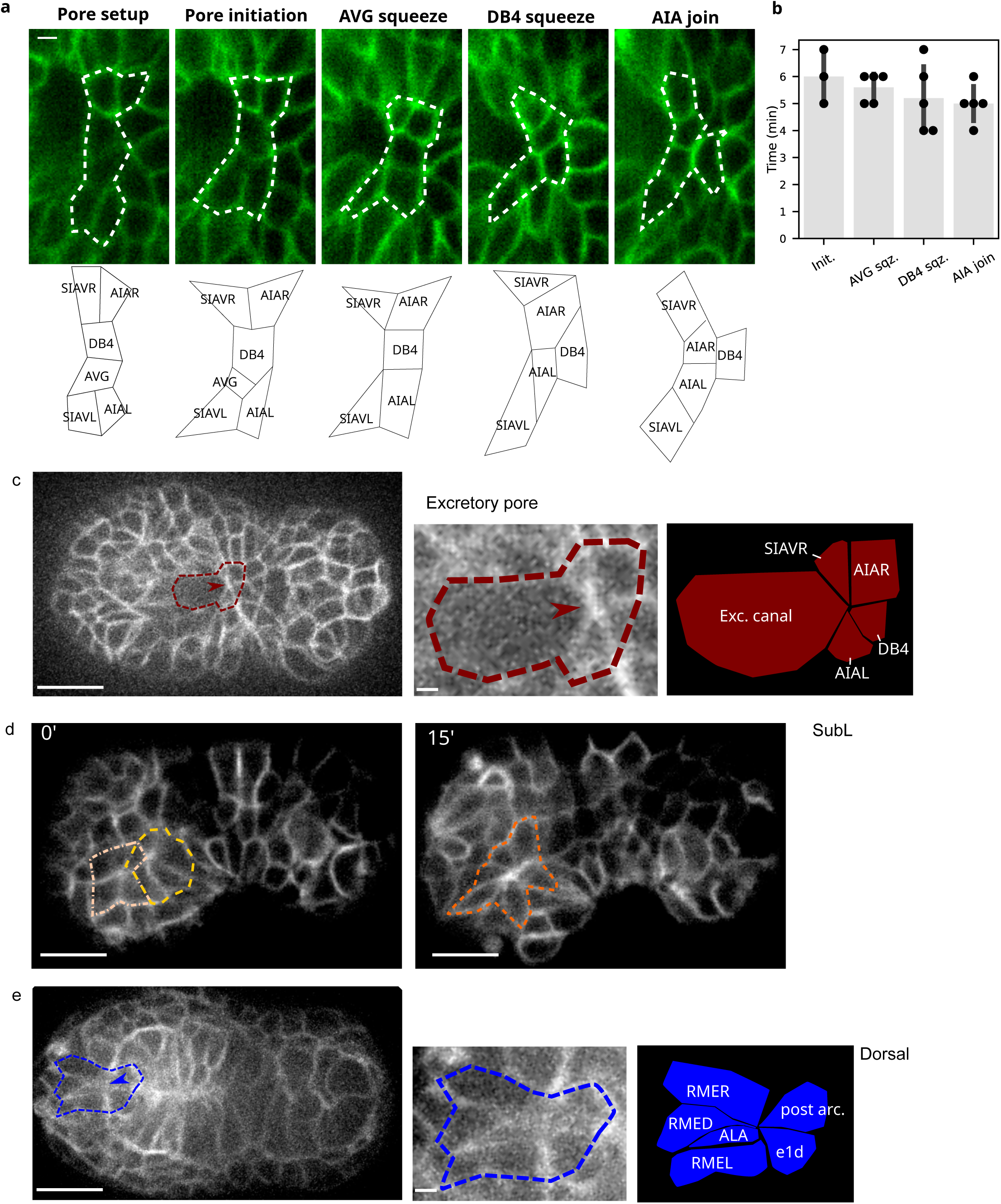
Lineage-based identification of nerve ring rosette cells. **a**, The ventral portion of the Exc. pore follows a 5-step stereotyped reference sequence of cell movements (AIAL, AIAR, AVG, DB4, SIAVL, SIAVR) beginning before and ending after the formation of the 8 rosettes shown in Fig. S1a. Top: illustrative cell sequence. White dashed line: outline of the six cells. Bottom: schematic illustrating cell movements. Steps: (i) Pore setup: cells align apposed to the excretory cell; (ii) Pore initiation: anterior-facing side becomes slightly curved; (iii) AVG squeeze: SIAVL and DB4 contact, pushing AVG ventrally, while the Exc. pore rosette converges to a central vertex; (iv) DB4 squeeze: AIAL and AIAR contact, pushing DB4 posteriorly; (v) AIA join: increased contact between AIAL and AIAR, anterior side of cells stiffens. Scale bar: 1 μm. **b**, Time (minutes) between subsequent stages in the Exc. pore sequence. For example, Pore initiation (Init.) is measured relative to Pore setup; AVG squeeze (sqz.) is measured relative to Pore initiation, etc. Bar height: average time. Error bars: standard deviation. Dots: measurements from individual embryos. **c**, Excretory pore rosette. Left: whole-embryo view with rosette outlined by a dashed line. Center: zoomed-in view of the outlined rosette. Right: identification of individual rosette cells. **d**, SubL rosette formation. Two smaller rosettes merge over 15 minutes to form the mature SubL rosette. **e**, Dorsal rosette formation. Left: whole-embryo view with rosette outlined by a dashed line. Center: zoomed-in view of the outlined rosette. Right: identification of individual rosette cells. **c**-**e**, Rosette cells were identified through lineage tracking from the 4-cell stage (Methods). ProxL and DistL rosettes are shown in Fig. 2b,e and are not included here. Scale bars: whole-embryo images: 10 μm; zoomed-in views: 1 μm.

**Figure S3:**
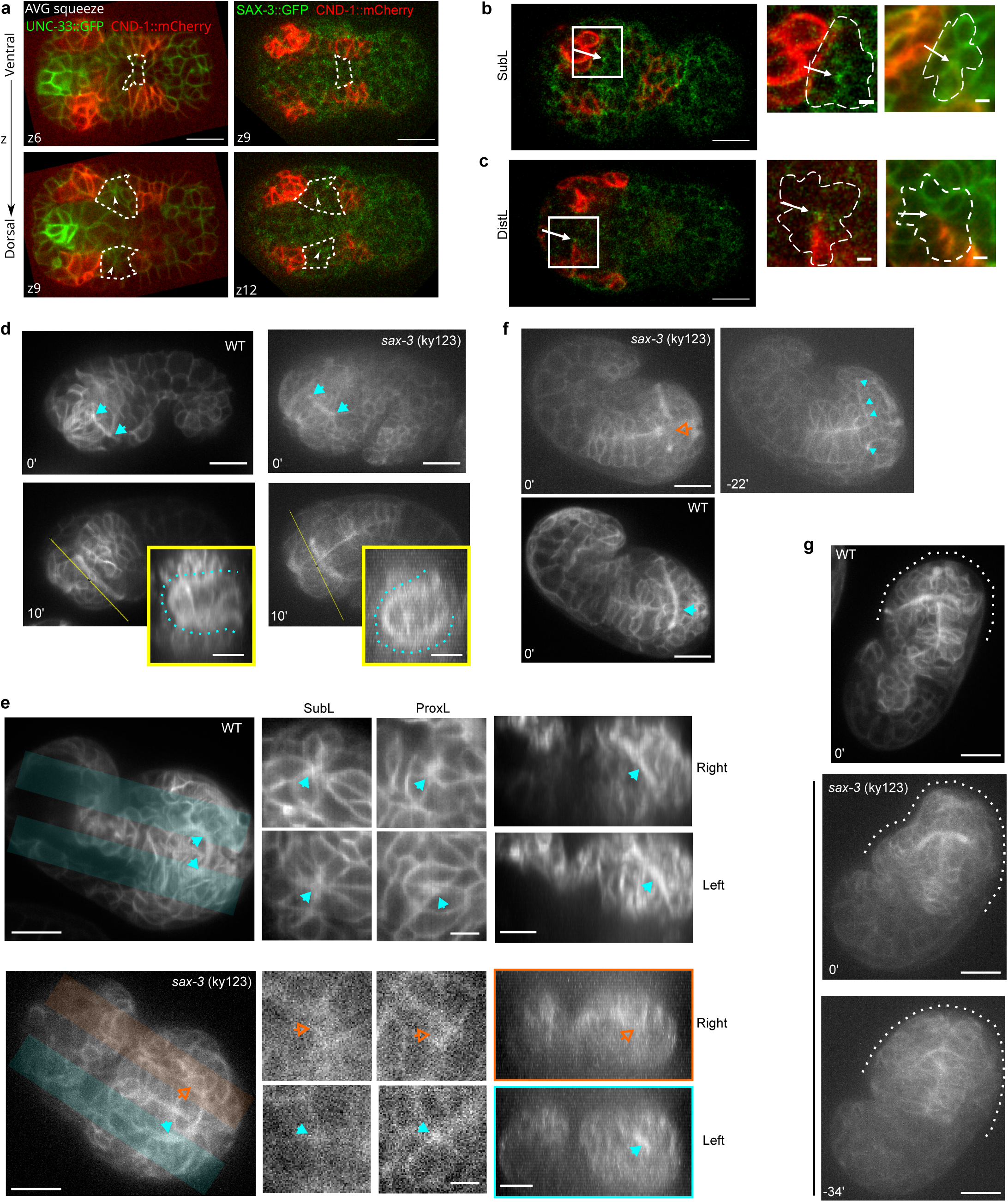
a-c, SAX-3 localizes to rosette centers. **a**, *cnd-1p*::PH::mCherry-expressing cells serve as spatiotemporal landmarks to locate rosettes. Left: UNC-33::GFP, CND-1::mCherry. Right: SAX-3::GFP, CND-1::mCherry. Both embryos aligned to the “AVG squeeze” stage (Fig. S1b). Dashed lines: contour of Excretory pore sequence cells (both) and, moving 3 Z-sections dorsally, the posterior SubL rosette (between the two CND-1::mCherry patches). Arrowhead: rosette center (left) and SAX-3 localization (right). **b**, SubL and **c**, DistL rosettes at a slightly later time-point. White rectangles indicate zoomed regions (right), labeled with a membrane marker. Arrows: rosette centers. **a**–**c**, Ventral view, anterior left. **d-g, Phenotypic consequences of *sax-3* loss are variable and dissociable.** Cyan filled / orange open arrowheads indicate presence/absence of the indicated structure throughout. **d, Incomplete penetrance: *sax-3* embryos range from WT-like to severely affected.** Fascicle outgrowth from SubL to ProxL (top) and nerve ring closure via orthogonal projection (bottom, yellow lines; dotted line marks closure), wild type (left) vs. *sax-3* (right), same embryos 10 min apart. **e, Rosette convergence is important for fascicle formation.** Rosette integrity corresponds to fascicle outgrowth: ventral view (left) with orthogonal validation of SubL/ProxL vertex presence on the right vs. left side of the same embryo, 15 min prior (middle), and maximum orthogonal projections confirming SubL fascicle status for the corresponding regions (right; top: WT, bottom: *sax-3*). **f, Rosette convergence alone is not sufficient for correct fascicle orientation.** Fascicle absence despite normal rosettes: *sax-3* embryo lacking fascicles across the dorsal midline (top left), the same embryo 22 min earlier showing left-side SubL/ProxL/DistL and right-side ProxL rosettes (top right), and a WT embryo with a fascicle crossing the midline for comparison (bottom). **g, Head morphology defects were excluded from scoring.** Head morphology defects: WT (top) and *sax-3* (middle) at a matched timepoint, and the same *sax-3* embryo 34 min earlier with normal head morphology (bottom); dotted lines mark head contour. WT embryos used an extrachromosomal marker; *sax-3* embryos used an integrated marker (Methods). WT and *sax-3* images shown together are distinct individuals, paired by matched embryo orientation. Scale bars: **a**, 10 µm; **b**,**c**, 1 µm; **d**–**g**, 10 µm except **e** middle, 2 µm.

**Figure S4:**
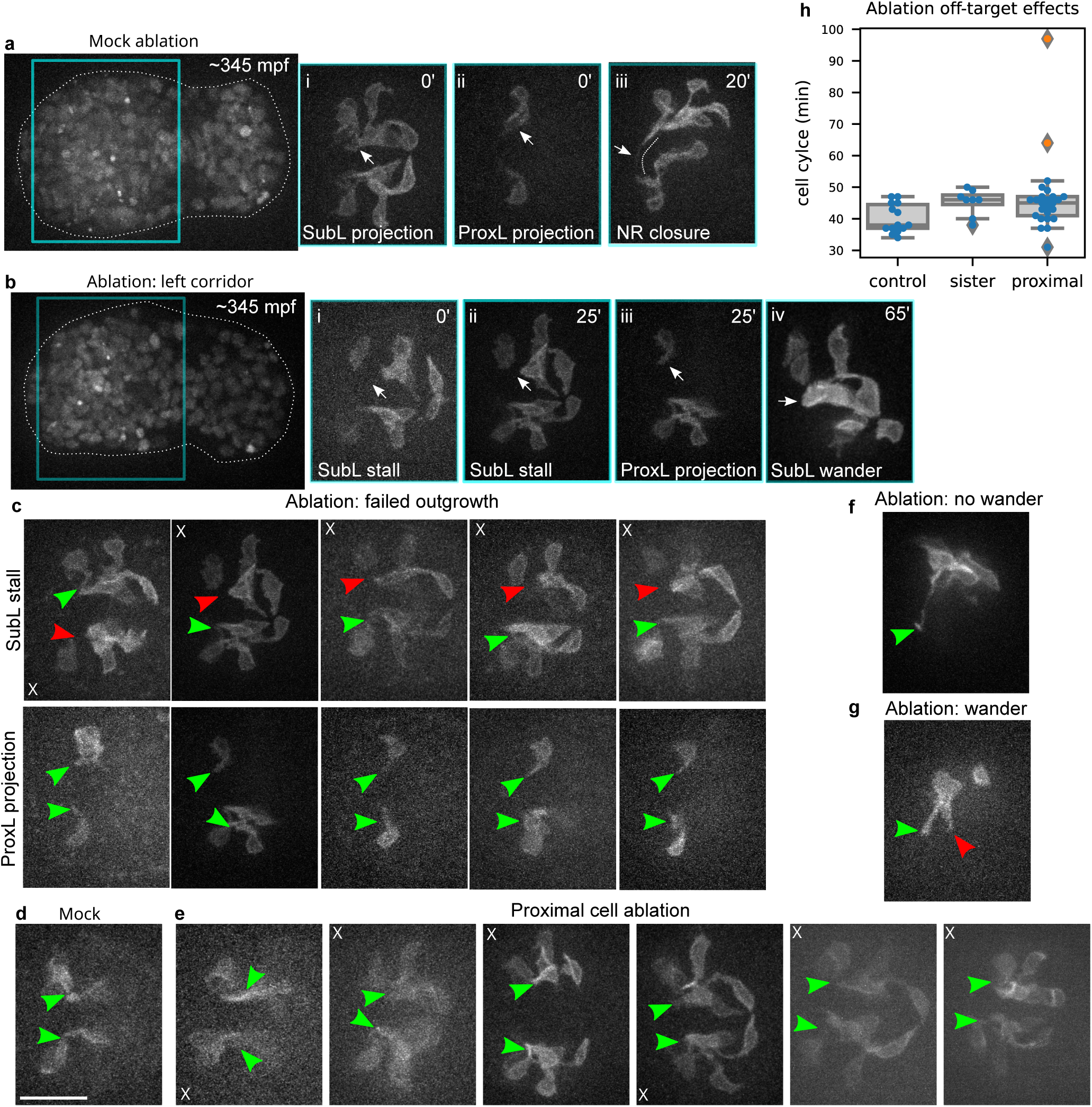
Specificity and penetrance of bridge cell ablation effects. **a**, Mock ablation control. Left: whole embryo (PIE-1::mCherry, nuclear marker for ablation targeting; LIM-4::GFP, SubL/ProxL outgrowth) staged at 345 mpf; cyan box marks the region shown in (i–iii), with time-points marked on each image. (i) SubL and (ii) ProxL projections at 0 min, different z-planes; (iii) nerve ring closure at 20 min (dashed line). **b**, Left corridor cell ablation. Left: whole embryo (same markers and staging as a); cyan box marks the region shown in (i–iv), with timepoints marked on each image. (i) SubL projection stalled at 0 min; (ii) SubL remains stalled at 25 min; (iii) ProxL projection at 25 min, different z-plane from (ii); (iv) SubL wanders outside the nerve ring path at 65 min. Apparent side-switching of SubL cell bodies reflects rotation of whole embryo during imaging. **c**, Ablation of the target corridor cell across all 6 successful ablations (columns), each showing SubL outgrowth failing to reach the ProxL rosette (top row) with confirmed ProxL projection (bottom row). **d**, Example mock ablation. **e**, Ablation of a cell proximal to the target corridor cell, showing normal SubL/ProxL outgrowth, as in mock. **f**, SubL stall followed by recovery of correct outgrowth — the one embryo, of five with initial stalling, that did not proceed to wander. **g**, In one embryo, wander was restricted to a single pioneer within the fascicle, while others in the same fascicle grew correctly — illustrating that the wander phenotype can be sub-fascicular rather than uniform across an embryo’s pioneers. **c**-**g**, Arrow heads indicate normal (green) or perturbed (red) phenotype. **h**, Evaluation of off-target effects on neighboring cells. Each dot represents a cell, with the y-axis showing cell cycle length (minutes) measured from 3d time-lapse imaging. Categories: Control = contralateral cells born at the same time as the ablated cell; Sister = sister cell of the ablation target; Proximal = adjacent cells, including those in the laser light path. Orange dots indicate outliers with prolonged cell cycles. Box plots show median (center line), upper and lower quartiles (box limits), whiskers = 1.5× interquartile range, and individual outliers. Two embryos exhibiting off-target effects were excluded from further analysis.

**Figure S5:**
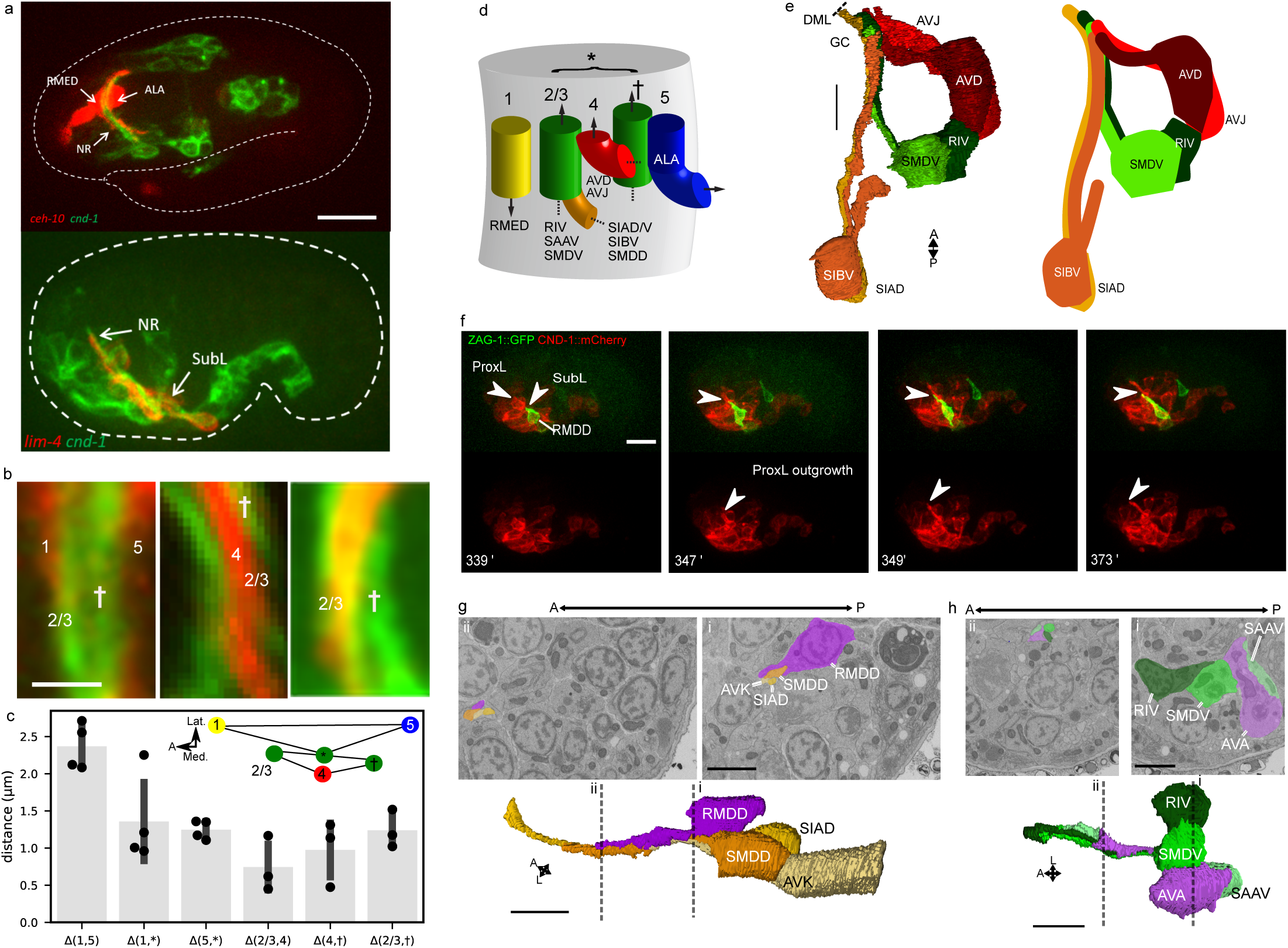
Triangulation of pioneer fascicle placement from fluorescence imaging. **a**, Fluorescence images of embryos with rosette axons labeled. Dashed white lines indicate embryo contour. Left: 2-fold embryo with *ceh-10p*::GFP and *cnd-1p*::PH::mCherry. Right: comma-stage embryo with *lim-4p*::GFP and *cnd-1p*::PH::mCherry. GFP and mCherry channels are recolored (red and green, respectively) for consistency with Fig. 4b. **b**, Relative positions of rosette axons. Bundles are labeled by numbers and † as in panel D. Left: *ceh-10p*::GFP, *cnd-1p*::PH::mCherry. Middle: *zag-1p*::PH::GFP, *cnd-1p*::PH::mCherry. Right: *lim-4p*::GFP, *cnd-1p*::PH::mCherry. **c**, Distances between fluorescent bundles labeled in panel **b** were measured across multiple embryos (Table S2, ‘pioneer_triangulation’). For example, Δ(1,5) is the distance between bundle 1 and 5. When comparing *ceh-10* and *cnd-1* expression, bundles 2/3 and † cannot be differentiated, so the distance from the center of *cnd-1* (*) to bundles 1 and 5 is used. Bar height: mean distance; error bars: s.d.; dots: individual measurements. All distances: n ≤ 4 embryos (limited by embryos with optimal orientation for visually discerning marker overlap). Inset: schematic showing triangulation of bundle positions, where nodes are bundles and line lengths represent average distances. **d**, Schematic of early pioneer fascicle hypothesis-consistent placement in the nascent nerve ring, triangulated from two-color fluorescence imaging. Arrows indicate direction of axon growth. SubL axons (orange) converge with ProxL axons (green). At later stages, current markers do not distinguish the two bundles. †: unidentified axon bundle. **e**, EM reconstruction of pioneer fascicle outgrowths in 365 mpf embryo. For clarity, only the two pioneers with the longest processes for each rosette are shown. Dashed line marks the dorsal midline (DML). GC: Growth cone. **f**, Timelapse imaging of RMDD (ZAG-1::GFP) outgrowth along SubL and ProxL fascicles (CND-1::mCherry). RMDD initially projects along SubL fascicles (0 minutes) but then tracks the ProxL fascicle at the ProxL rosette point (24 minutes). **g,h**, Top: EM sections showing proximal (i) and distal (ii) segments of pioneer neurites; bottom: volumetric reconstructions of the same segments (dashed lines, i and ii). **g**, RMDD and SubL (SIAD, SMDD, AVK). **h**, AVA and ProxL (RIV, SMDV, SAAV). No fluorescent embryonic marker was available for AVA at the time this dataset was collected, as EM data motivating its inclusion as a follower of interest was not yet available; RMDD is supported by both fluorescence tracking and EM reconstruction, which converge on the same conclusion: the fluorescence-observed tracking corresponds to genuine ultrastructural contact, ruling out an unmarked intervening cell, while the EM-confirmed contact arises from active tracking rather than passive convergence. This bidirectional agreement supports extending each single-modality inference to the evidence available for AIY (tracking behavior, fluorescence only) and AVA (genuine contact, EM only). **c**, A: anterior; Lat: lateral; Med: medial. **g,h**, A: anterior; L: left. Scale bars: **a**, 10 μm; **b**, 1 μm; **e,g,h**, 2 μm.

**Figure S6:**
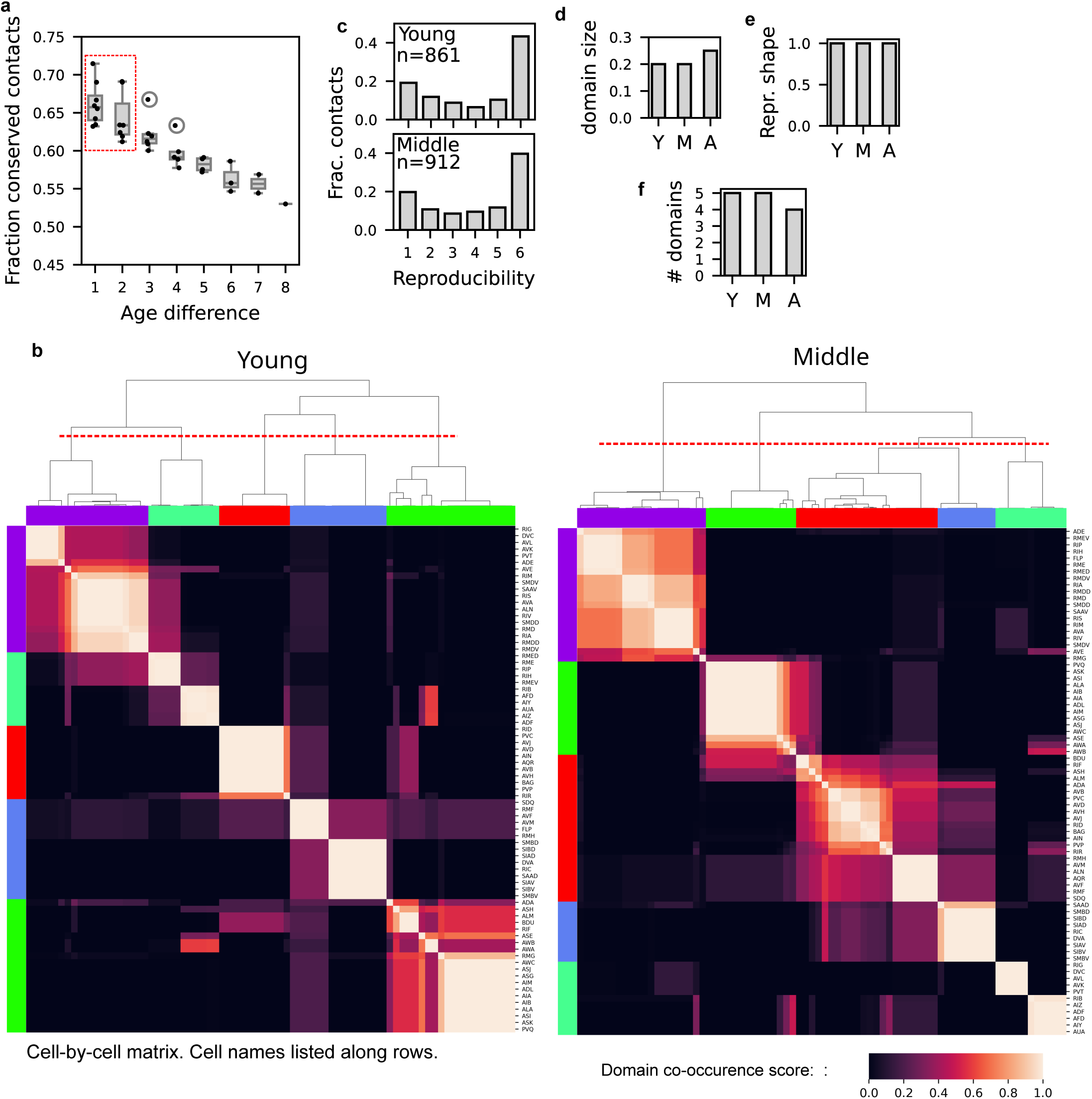
Age-independent structural features of nerve ring organization. **a**, Fraction of membrane contacts conserved between EM datasets (y-axis) as a function of age difference (x-axis). Pairwise age differences are based on the chronological ordering of datasets. Individual dots represent pairwise comparisons. Boxes show the range of conserved fractions, and the bar indicates the median. Red dashed box highlights that age differences less than 3 are most similar and can be grouped together for reproducibility analysis. **b**, Spatial domain partitioning of the most reproducible contacts in the young and medium age groups. Matrix entries show the probability that a pair of cells occupy the same spatial domain (0–1) when using a population-based clustering algorithm (Methods). Red dashed line: cut-off for hierarchical clustering assignment of the dendrogram. Color bars indicate neuron spatial domain assignment. **c**, Reproducibility of axon contacts across 6 datasets in the young and medium age groups. y-axis: fraction of pairwise axon contacts; x-axis: number of datasets where the pairwise contact occurs. Bimodal reproducibility is quantified as the distance between the two highest distribution peaks. n: total number of pairwise axon contacts used in the calculation. **d**, Bimodal reproducibility across the 3 age groups. **e**, Number of spatial domains across the 3 age groups. **f**, Average domain sizes across the 3 age groups. **d**–**f**, These values define the empirical benchmarks used for SDM calculations. Age group assignments. Y: young. M: medium. A: adult.

**Figure S7:**
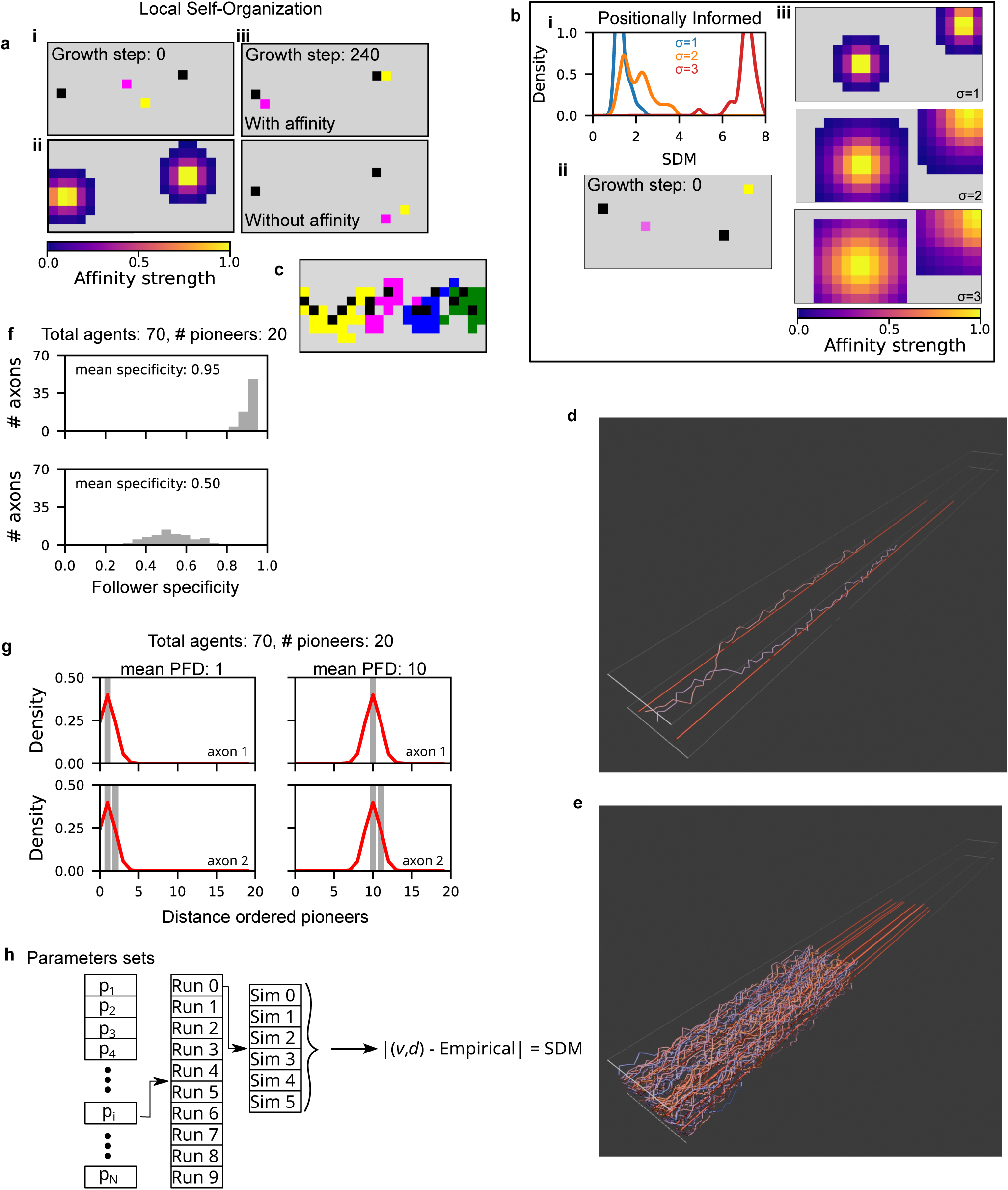
Agent-based simulation framework for nerve ring innervation. **a**, Overview of local self-organizing simulation models: Gray simulation boxes show transverse planes along simulation volume. **a**i, At initialization, fixed pioneer agents (black) and non-fixed follower agents (colored) are randomly placed within an inner region of the grid (dashed red box). **a**ii, When pioneer–follower affinity is enabled, pioneers generate an attractive Gaussian field (peak = 1, standard deviation = 1 grid unit). **a**iii, With affinity enabled, followers innervate toward their matched pioneers by the end of growth (240 steps); when affinity is disabled, followers innervate toward random positions **b**, Overview of positionally informed simulation models: Baseline control. **b**i, Kernel density estimates of average simulation distance metric (SDM) values for PI simulations at increasing positional uncertainty (σ = 1, 2, 3). **b**ii, At initialization, pioneer and follower agents are placed within the grid without pioneer–follower affinity. **b**iii, Follower trajectories are biased by an attractive Gaussian field, with positional uncertainty, represented by σ, controlling the strength of the bias toward the target position **c**, Example spatial domain organization with pioneer–follower affinity enabled (14 pioneers, 55 followers). Followers are colored by spatial domain assignment. Shown for illustration only; quantitative comparisons are in Fig. 6b,f. **d**, 3d reconstruction of a simulation with two pioneers (red tubes) and two followers (multi-color tubes). White box indicates simulation volume (Movie S6). **e**, 3d reconstruction of a simulation with 14 pioneers and 56 followers (Movie S7). **f**, Follower specificity is randomly assigned for each follower using a binomial distribution. Shown are example specificity distributions for mean specificity 0.95 (top) and 0.5 (bottom), with 70 axons and 20 pioneers. **g**, Pioneer–follower distance is drawn from a Gaussian distribution. Plots show the probability density (red) of ranked follower–pioneer distances (x-axis). Left and right panels show mean distances of 1 and 10, respectively; top and bottom panels show one or two selected pioneers, as indicated. When specificity is low, multiple selected pioneers tend to be spatially clustered. Pioneer distinctiveness and pioneer–pioneer distance are assigned analogously. **h**, Parameter sampling strategy. For each experiment, many parameter combinations are sampled. For each combination, multiple runs are initialized from different agent placements, with six simulations per placement. Summary statistics for reproducibility shape (v) and spatial domains (d) are computed. The simulation distance metric (SDM) quantifies deviation from empirical values.

**Figure S8:**
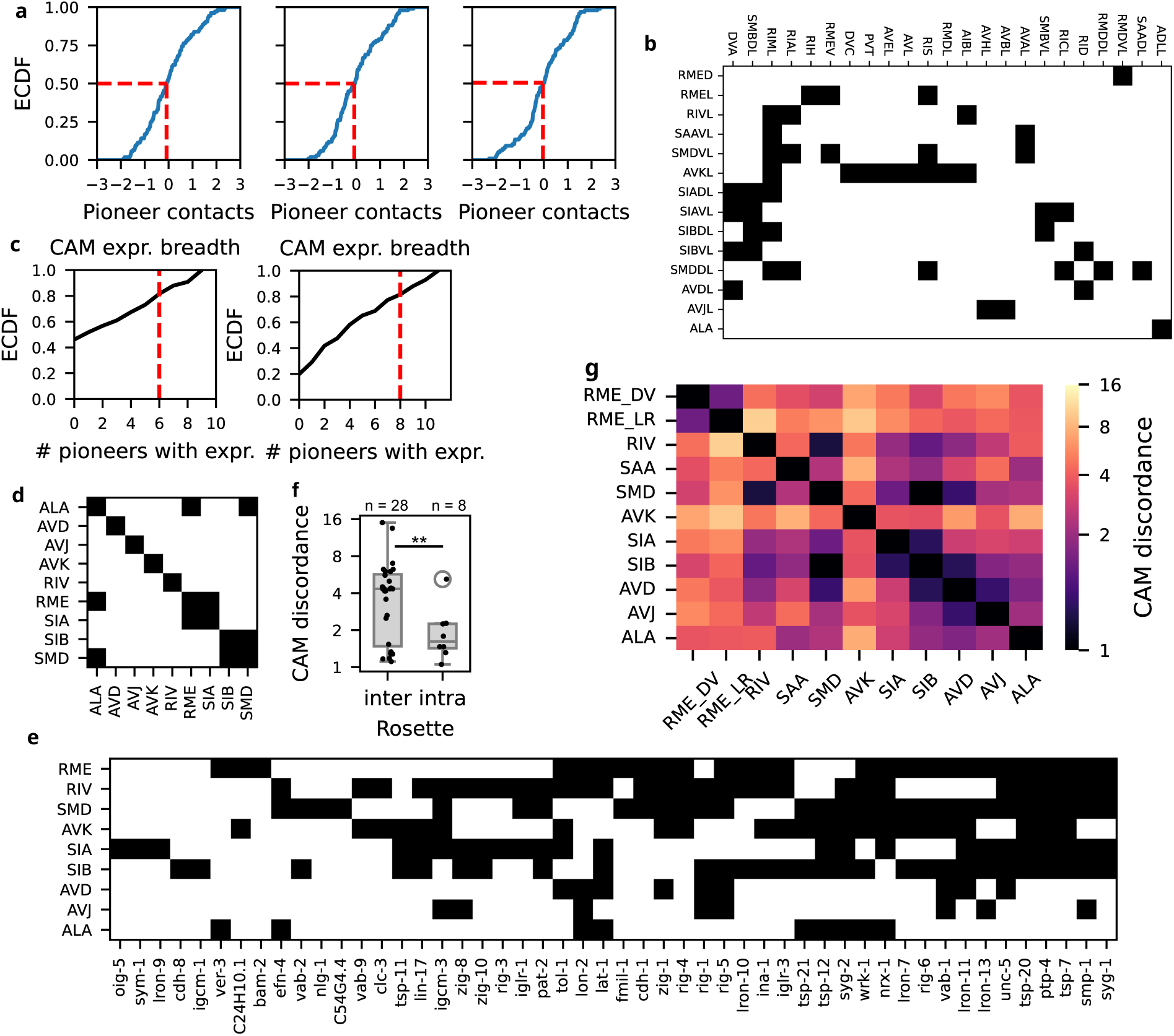
Defining shared-pioneer contact and molecular distinctiveness using CAM expression. **a**, Cumulative distribution of pairwise surface area contacts between axons for the young, middle, and adult age groups. Pairs with above-average contact (red dashed line) are used to define shared-pioneer contact (see Methods). **b** Follower (rows) by pioneer (columns) table showing which followers share above-threshold conserved contact with which pioneers; underlying data for the domain-membership analysis in Fig. 5g. **c**, Cumulative distribution of the number of pioneers expressing each CAM in the embryo (left) and L4 (right). Only CAMs expressed in fewer than 75% of cells are included (red line). **d**, Pioneer similarity table based on embryonic single-cell data. Pioneer pairs with low CAM discordance are defined as similar (black entry). A pioneer is considered more distinctive when it shares fewer CAMs with other pioneers. **e**, Pioneer-by-gene matrix for the embryo. Black entries indicate the presence of CAM expression in that pioneer. **f**, Boxplot of pairwise CAM discordance between pioneers within (intra) and from different (inter) rosettes, computed from the embryonic expression data in e. Boxes show median and interquartile range; dots show individual pairwise pioneer contacts. Welch’s t-test; *p*=0.013. **g**, CAM discordance heatmap for the L4. Intensity indicates the level of CAM discordance between pioneer pairs.

**Figure S9:**
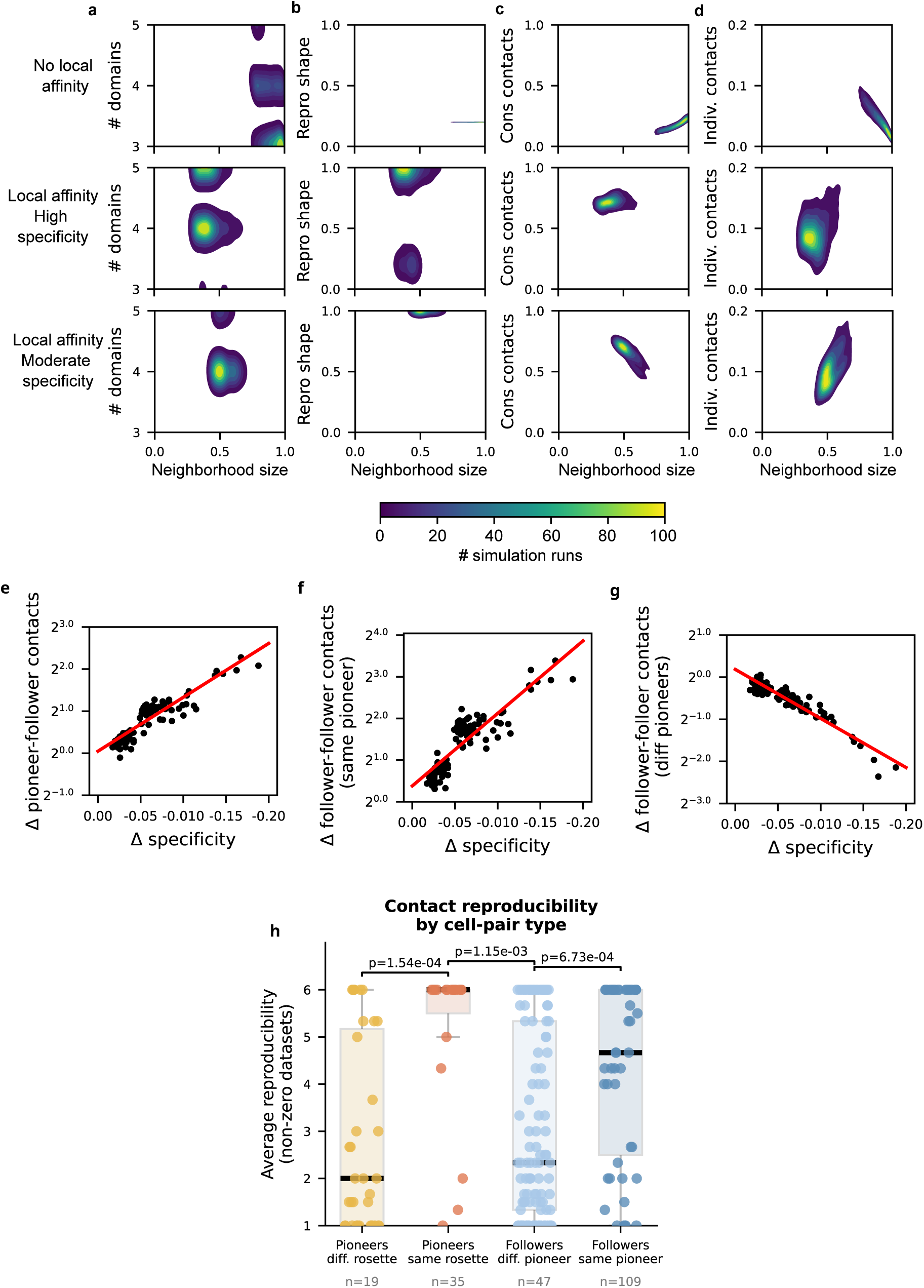
Constrained neighborhood size and reduced specificity increase reproducibility. **a**, 2d density plot of number of spatial domains versus average neighborhood size. **b**, 2d density plot of reproducibility shape versus average neighborhood size. **c**, 2d density plot of fraction of conserved contacts versus average neighborhood size. **d**, 2d density plot of fraction of individual contacts versus average neighborhood size. **a**-**d**, Top: Simulations without affinity. Middle: Simulations with affinity and high (>90%) follower specificity. Bottom: Simulations with affinity and moderate (80-90%) specificity. Density intensity is the number of simulations. **e**, log_2_ fold change in the fraction of conserved pioneer–follower contacts as a function of follower specificity. **f**, log_2_ fold change in the fraction of conserved follower–follower contacts as a function of follower specificity, where followers share at least one pioneer in common. **g**, log_2_ fold change in the fraction of conserved follower–follower contacts as a function of follower specificity, where followers share no pioneers in common. **e**–**g**, The x-axis shows the decrease in specificity from high to moderate; the y-axis shows the log_2_ ratio (moderate/high) of conserved contacts when decreasing the specificity. Each point compares matched simulations run at high and moderate specificity using identical initial configurations. Red line, linear fit. *n*=108 independent paired specificity comparisons. **h**. Distribution of reproducibility across contact types: pioneers in same (*n*=19) or different (*n*=35) rosettes (*p* = 1.54 × 10^−4^); followers with shared (*n*=47) or different (*n*=107) pioneers (*p* = 6.73 × 10^−4^). The pioneer-level comparison is included for completeness; the text focuses on the follower-level result, which more directly bears on the specificity/neighborhood relationship discussed in this section. Individual points represent pairwise contacts; boxplots show the median and interquartile range, with whiskers extending to the observed range excluding outliers. *p*-values: Mann Whitney test.

